# Genetic deconvolution of embryonic and maternal cell-free DNA in spent culture medium of human preimplantation embryos through deep learning

**DOI:** 10.1101/2025.01.20.633979

**Authors:** Zhenyi Zhang, Jie Qiao, Yidong Chen, Peijie Zhou

## Abstract

Noninvasive preimplantation genetic testing for aneuploidy based on embryonic cell-free DNA (cfDNA) released in spent embryo culture media (SECM) has brought hope in selecting embryos that are most likely to implant and grow into healthy babies during assisted reproduction. However, maternal DNA contamination in SECM significantly hampers the reliability of embryonic chromosome ploidy profiles, leading to false negative results, particularly at high contamination levels. Here, we present DECENT (<u>de</u>ep <u>c</u>opy number variation (CNV) r<u>e</u>co<u>n</u>struction), a deep learning method to reconstruct embryonic CNVs and mitigate maternal contamination in SECM from single-cell methylation sequencing of cfDNA. DECENT integrates sequence features and methylation patterns by combining convolution modules, long-short memory, and attention mechanisms to infer the origin of cfDNA reads. The benchmarking study demonstrated DECENT’s ability to estimate contamination proportions and restore embryonic chromosome aneuploidies in samples with varying contamination levels. In contaminated SECM clinical samples, including one with more than 80% maternal reads, DECENT achieved consistent CNV recovery with invasive tests. Overall, DECENT contributes to enhancing the diagnostic accuracy and effectiveness of cfDNA-based noninvasive preimplantation genetic testing, establishing a robust groundwork for its extensive clinical utilization in the field of reproductive medicine.

## Main

The frequency of embryo aneuploidy increases exponentially with advancing maternal age and is a major cause of pregnancy failure, miscarriage, and congenital anomalies in both natural conception and in vitro fertilization (IVF)^1–4^. Reproductive medicine faces a significant challenge in identifying embryos with the highest potential for successful live births. Preimplantation genetic testing for aneuploidies (PGT-A) has emerged as a vital tool for assessing chromosome abnormalities, particularly given their frequent occurrence in human embryos, challenging conventional morphological assessments alone. Early randomized trials on the clinical utility of PGT-A have been controversial. Some studies have suggested that PGT-A improves pregnancy rates or implantation rates, while other studies have failed to prove the clinical benefits of PGT-A^5–10^. A recent systematic review of 11 randomized trials revealed that PGT-A has a significant effect on improving live birth rates in advanced-stage pregnant women (>35 years old)^11^. While several techniques for PGT-A exist, including biopsy methods such as polar body, blastomere, or trophectoderm (TE) biopsies, TE biopsy has garnered increasing interest for its efficacy. However, inherent limitations, such as convenience, invasiveness, potential misdiagnosis due to mosaicism, and concerns regarding impacts on implantation potential, underscore the need for alternative approaches.

Recent advancements have explored noninvasive preimplantation genetic testing of aneuploidy (niPGT-A) using cell-free DNA (cfDNA) extracted from spent embryo culture media (SECM), offering a promising avenue to circumvent the limitations of traditional biopsy methods. Some studies have compared the ploidy concordance of SECM-based noninvasive PGT-A with that of TE biopsies or ICM cells, revealing that the potential of cfDNA for indicating embryo ploidy may be superior to that of TE biopsies^12,13^. However, challenges persist due to maternal DNA contamination in SECM, leading to gender discordant and false negative results. A single-nucleotide polymorphism (SNP) study revealed greater maternal DNA contamination (86-94%) in SECM^14^.

Methylation patterns play a pivotal role in embryonic development and distinguishing cfDNA cell types^15–18^, tracing their origins across diverse contexts, including disease detection and diagnosis^19^, prenatal testing, and organ transplant monitoring^20^. Moreover, methylation markers can also be utilized to screen for high-quality embryos, distinguishing them from those of lower quality^21^. Our previous study used whole-genome DNA methylation sequencing to identify cfDNA in culture media originating from blastocysts, cumulus cells, and polar bodies. Severe maternal contamination (>60%) was present in one-third of SECMs. Specifically, the gender discordant and false negative rate of niPGT-A increased with maternal contamination. Once maternal contamination exceeds 60%, the embryo chromosome ploidy inferred by SECM is not accurate, highlighting the critical nature of addressing maternal contamination, as higher contamination ratios significantly compromise the reliability of chromosome ploidy profiles^22,23^.

Existing strategies to reduce contamination levels primarily concentrate on adjustments in the sample collection process. For instance, altering the SECM collection time, embryo rinsing protocols, culture media renewal, and gentle re-denudation of residual cumulus cells. An extended culture duration results in a greater yield of cfDNA present in the culture medium. SECMs from embryos cultured until day 6/7 exhibit greater informativeness and consistency than those from embryos cultured until day 5^22,24^. Compared to the one-step embryo rinsing protocol, the sequential method showed markedly superior performance^25^. Adding extra medium renewal during day 4 embryo culture enhances the agreement of niPGT with TE biopsy^26^. Re-denudation of residual cumulus cells on day 3 decreases the impact of maternal contamination and enhances the precision of cfDNA detection in SECM^27^. However, these methods have limited efficacy in eliminating maternal contamination and require additional effort, increasing the workload of IVF laboratory personnel. Therefore, there is an urgent need to develop simpler and more efficient methods to remove maternal contamination.

In recent years, deep learning models, particularly those inspired by natural language processing (NLP) tasks, have emerged as powerful tools for analyzing sequence-like data in biological research^28–34^. Within the genomics domain, various deep learning frameworks, including recurrent neural networks (RNNs), long short-term memory networks (LSTMs), and transformers, have been successfully applied to tasks such as transcription factor binding, mutation detection, chromatin accessibility assessment, and promoter/enhancer region identification^29,35–38^. Moreover, in the analysis of cfDNA, particularly methylation data, deep learning methods have demonstrated efficacy in disease diagnosis and prediction, as well as monitoring treatment outcomes^19,39–47^. For example, these methods have been employed for tissue deconvolution from plasma cfDNA to aid in cancer diagnosis and early screening, as well as for monitoring treatment side effects. Despite these advancements, comparable algorithm designs and research on embryonic and maternal cfDNA are still lacking. It remains unclear whether deep learning methods could effectively address the challenges such as cfDNA deconvolution and embryonic CNV reconstruction posed by maternal DNA contamination in SECM.

Here, we present deep CNV reconstruction (DECENT), a new deep learning framework aimed at mitigating maternal contamination in SECM and reconstructing embryonic copy number variations (CNVs). DECENT leverages sequence and methylation information from both embryonic and maternal sources, utilizing convolutional neural networks and attention mechanisms to infer the origin of sequence reads. The key features of our method include the following: 1) the use of a substantial dataset (∼15 M) encompassing both embryonic and maternal sequence and methylation features, 2) the use of a new method for estimating maternal contamination proportions in SECM, and 3) the ability to remove maternal contamination and reconstruct embryonic copy number variations. We systematically benchmarked the ability of DECENT to infer maternal contamination levels and reduce false negatives of chromosome aneuploidy analysis in simulated SECM samples of various contamination levels. CNV analysis of 194 real clinical SECM samples was then performed with DECENT. We also applied interpretable machine learning tenues to the DECENT classifier to elucidate the potential differential methylation patterns between embryonic and maternal cfDNA reads.

## Results

### Overview of DECENT

DECENT aims to accurately reconstruct chromosome ploidy profiles of embryonic cfDNA in SECM for niPGT-A. A significant challenge is the presence of moderate to severe contamination of cumulus cell DNA in SECM. Therefore, precise differentiation and extraction of cfDNA from embryos for CNV calculation in SECM is crucial for the clinical use of noninvasive cfDNA-based approaches.

To ensure the reliability of CNV detection, we first employed a deep neural network to identify the likelihood of each read originating from embryonic cfDNA (Methods). Both the sequence data and the corresponding methylation data obtained from the spent embryo media were utilized for training and validation (Fig. **1A**). Once these probability scores were obtained, we then determined a threshold that balances precision and the number of reads. Reads with a probability score below the specific threshold are identified as originating from the embryo and are subsequently used in chromosome CNV detection. This ensures that the reads used for CNV detection are predominantly embryonic, thereby increasing the reliability of CNV analysis. Furthermore, probability scores can also be used to compute the degree of cumulus cell contamination within the culture medium through maximum likelihood estimation. By comparing these estimates with those of other existing methods, we can assess the performance of DECENT and apply them to address the challenging CNV detection task in highly contaminated samples (Fig. **1B**).

**Fig 1.**
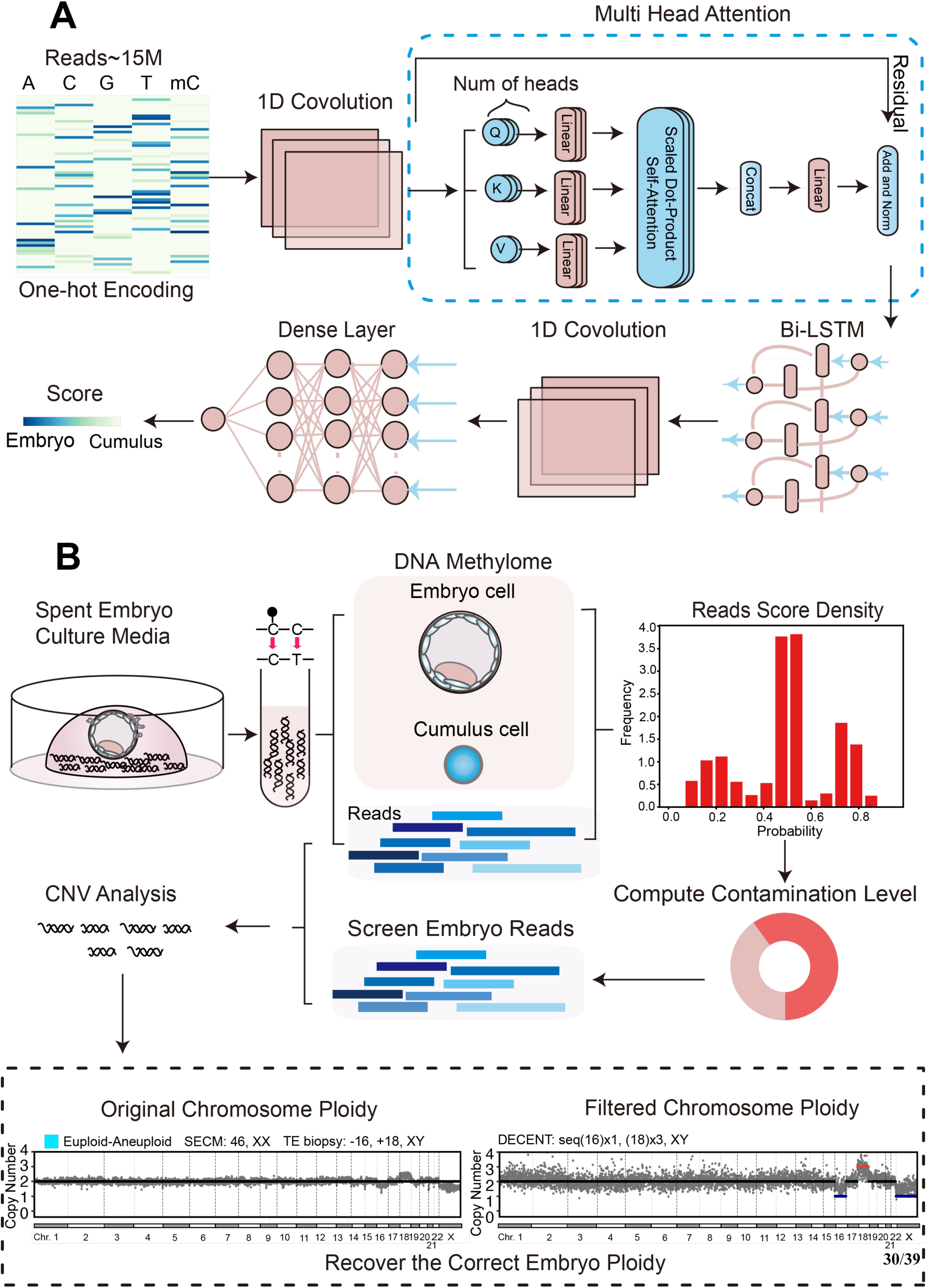
Overview of DECENT. **(A)** Schematic representation of the deep neural network employed for reads categorization. The network was designed to evaluate the likelihood of each read originating from embryonic cfDNA. **(B)** Schematic illustration of the procedure used to reconstruct embryonic chromosome ploidy contaminated by cumulus cells in SECM. Probability scores derived from the deep learning model are utilized to estimate the extent of cumulus cell contamination within the SECM. Subsequently, embryonic reads were filtered based on the established probability score threshold and retained for downstream analysis (sample n=194). In the CNV analysis, the light blue block denotes samples with euploid conditions in SECM and aneuploid conditions in TE.

### DECENT captures sequence features of cfDNA from embryo and cumulus cells

To evaluate the performance of the model, we first investigated the distribution of scores generated by the neural network model on the training dataset (Fig. **2A**). We observed a symmetric distribution of scores with three main modes concentrated at approximately 0.2, 0.5, and 0.7. The concentration of scores of approximately 0.2 and 0.7 suggested that the neural network successfully captured certain features specific to embryos and cumulus cells, enabling their discrimination. The clustering of approximately 0.5 may be attributed to a significant proportion of reads that can be present in both the embryo and cumulus categories, leading to the inability to distinguish such reads. However, this does not affect the efficacy of the method since, in the CNV detection task, precision is more critical than recall, as we aim to strictly control the selection of reads dominated by embryos to reconstruct embryonic CNVs. Therefore, we can only consider the high-confidence reads correctly classified as embryonic by the neural network rather than aiming for full coverage.

**Fig 2.**
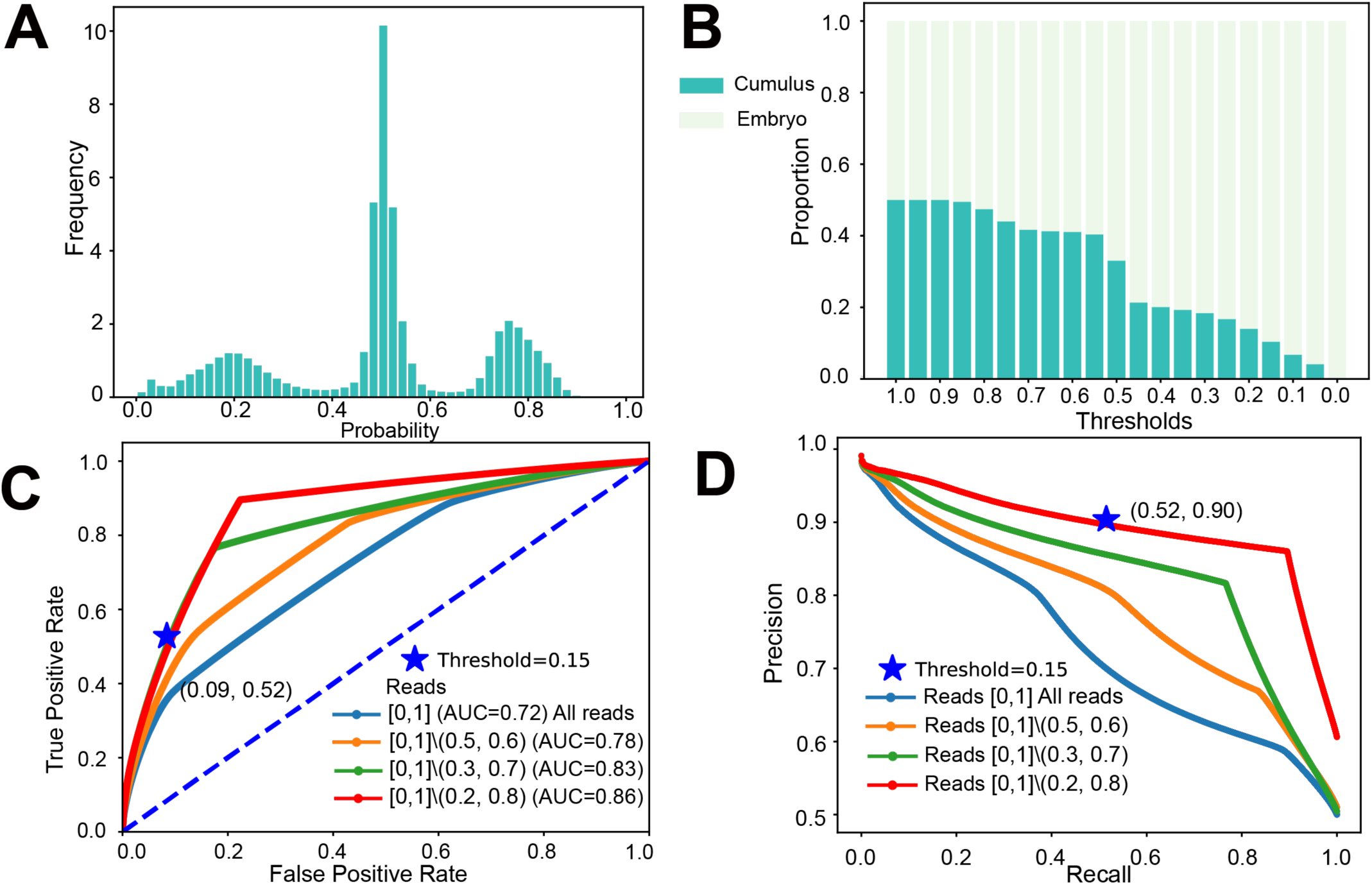
Evaluation of DECENT. **(A)** Distribution of scores generated by the neural network model on the training dataset, showing three main modes concentrated at approximately 0.2, 0.5, and 0.8. **(B)** Variations in the proportions of identified cumulus contaminants with different threshold values in the training dataset. A decrease in the proportion of cumulus contamination is observed as the threshold decreases. **(C)** Receiver operating characteristic (ROC) curves comparing the performance of the deep learning model under different threshold ranges for reads exclusion. Reads [a,b]\(c,d) means the reads whose score ranges in [a, c] and [d, b]. The pentagram marker indicates the value of the corresponding metric at threshold 0.15. **(D)** Precision‒recall (P‒R) curves demonstrate the trade-off between precision and recall for the deep learning model under different threshold ranges for read exclusion. Reads [a,b]\(c,d) means the reads whose score ranges in [a, c] and [d, b]. The pentagram marker indicates the value of the corresponding metric at threshold 0.15.

It is important to establish threshold values for calculating probability scores, defining reads below this score as embryos, which are subsequently used for downstream analysis (Methods). An appropriate threshold is essential because it balances the trade-off between retaining true embryo variations and minimizing noise introduced by cumulus contamination. A high threshold may lead to the inclusion of substantial cumulus contamination, obscuring real embryo variations and compromising the removal of false negatives. Conversely, a low threshold may result in the retention of too few reads, amplifying noise in subsequent CNV detection. We compared the variation in the proportion of cumulus contamination with different threshold values in the training dataset (Fig. **2B**). We observed a decrease in the proportion of cumulus contamination as the threshold decreased. Notably, selecting a threshold of 0.15 yielded a precision of approximately 90% based on the results from the training dataset, which may represent a favorable precision level.

We assessed the performance of the deep learning model using Receiver operating characteristics (ROC) and precision-recall curves (Methods). As previously mentioned, a portion of reads may be poorly classified by the deep learning model, resembling random classification. Since we are primarily interested in removing this portion of reads, we implemented various read score ranges that were strategically designed to facilitate the exclusion of undesirable reads while maintaining the integrity of embryonic sequences. In computation, we set the embryonic reads to be the positive class and the maternal reads to be the negative class to ensure better clarity. Through analysis, we observed an increase in the area under the curve (AUC) as intermediate score reads were progressively excluded. This increase in the AUC signifies an improved discrimination ability of the neural network in identifying embryonic reads amidst contamination (Fig. **2C**). Additionally, precision-recall curve analysis demonstrated an accompanying increase in recall for the same precision level, further suggesting the effectiveness of our approach (Fig. **2D**). Notably, at the final read scores range, our neural network achieved an AUC value of 0.86, which is indicative of its satisfactory performance in this task. Furthermore, we test the model’s classification performance by focusing on Differentially Methylated Regions (DMRs, Methods). We evaluated the impact of incorporating DMR-based enrichment prior to model training. We found that applying DMR enrichment resulted in a significant increase in AUC, achieving values up to 0.97 (Supplementary Fig. **S12 A, B**). However, DMR enrichment poses a substantial reduction in the number of reads retained. The diminished read count would impair our ability to detect CNVs reliably, which is a crucial aspect of our research. So, we do not adopt DMR during our training.

To further explore how our models learn the features of embryonic and maternal DNA and to assess the influence of different features on model predictions, we identified 6,259 reads from the training dataset, comprising typical maternal reads (high score) and typical embryonic reads (low score). These reads were processed through the model, and we extracted features from various layers. We reduced the dimensionality of the extracted features using Principal Component Analysis (PCA) and plotted their distribution in the first two principal components (Supplementary Fig. **S9**). We found that progression through various layers resulted in a more distinct aggregation of each class with a well-defined separation, underscoring the progressive enhancement of feature discriminability. These findings demonstrate the contribution of different network layers to the model’s predictive performance.

### DECENT estimates cumulus contamination levels in SECM

Estimating the proportion of maternal contamination within SECM samples is crucial for downstream analysis, as maternal contamination directly affects the accuracy of CNV. Following the application of DECENT to assign a score to each read, we utilized these scores to estimate the proportion of maternal contamination within the sample (Methods).

In the previous study^22^, the authors proposed a method for estimating the maternal contamination proportion using a DNA methylation signature. We compared the proportions computed using DECENT with those derived from methylation levels (Fig. **3A**, sample n=194). Notably, we observed a strong correlation between the proportions calculated by our deep learning method and those derived from methylation levels, indicating a high level of consistency. Despite the utilization of different methods, the consistency of the results cross-validates the effectiveness of our deep learning approach.

**Fig 3.**
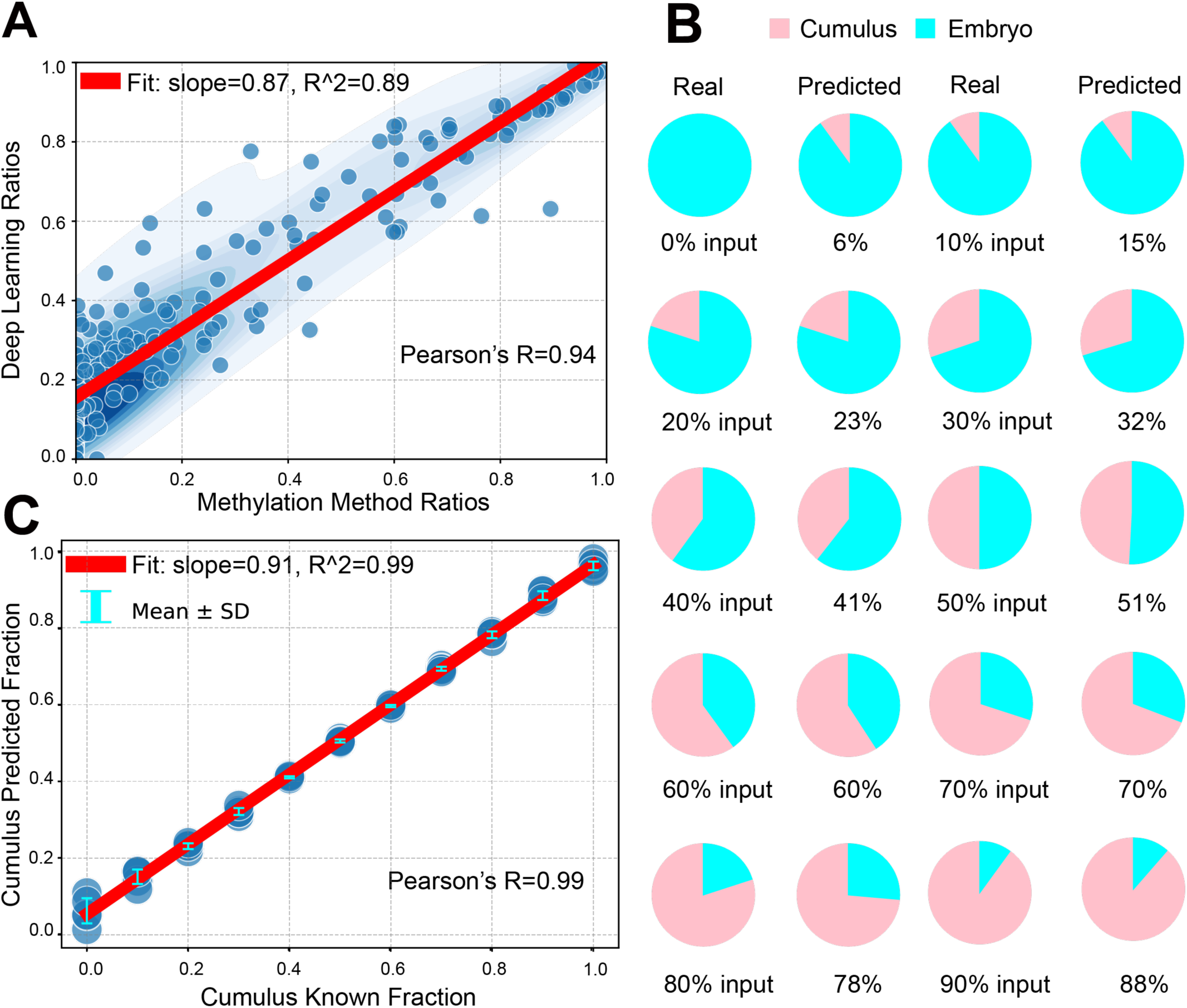
Validation of the estimated maternal contamination proportion by DECENT. **(A)** Estimation of maternal contamination proportion within SECM using our deep learning framework (sample n=194, P value < 0.001). **(B)** Simulation analysis of the estimated and input percentages of read mixtures containing varying proportions of embryonic and cumulus cell cfDNA. We show the mean estimated ratio across five independent runs. **(C)** Correlations between the predicted and input fractions of the simulated DNA mixtures (P value < 0.001).

To further verify the accuracy of DECENT in the simulation datasets, we generated a series of synthetic datasets with varying proportions of cumulus and embryo mixtures (Fig. **3B, C**, Methods). The estimated percentages exhibited a strong correlation with the input percentages of the DNA mixtures, as evidenced by linear regression lines (R = 0.99, Pearson’s correlation, t-test, Fig. **3C**). Overall, the simulation and benchmarking analysis support the utility of DECENT for quantifying the maternal contamination level of cfDNA in SECM samples.

### DECENT accurately reconstructs the embryonic chromosome aneuploidy in simulated contaminated samples

To evaluate the ability of DECENT to accurately depict embryonic chromosome aneuploidy, we reconstructed the chromosome aneuploidies using synthetic datasets mixed with different cumulus cell contamination (Methods). Initially, we selected an aneuploidy SECM sample (S53) with a contamination proportion of 0, representing reads exclusively derived from embryonic cfDNA, and the bulk cumulus sample (G4) was assumed to contain reads solely from cumulus cells. Subsequently, we introduced cumulus cell reads into the embryonic sample, simulating maternal contamination scenarios at contamination proportions of 60%, 65%, 70%, and 75% (Fig. **4A**). Our findings indicated that the introduction of contamination obscured the detection of original embryonic chromosome aneuploidies, such as the specific instances of seq(16)x1, (18)x3, and XY gender, with these variations being masked by maternal contamination. Moreover, as the contamination proportion increased, the degree of obscuration intensified. Notably, our algorithm successfully recovered at least one variation even at contamination proportions of 60%, 65%, 70%, and 75%, with gender correctly identified below 75%.

**Fig 4.**
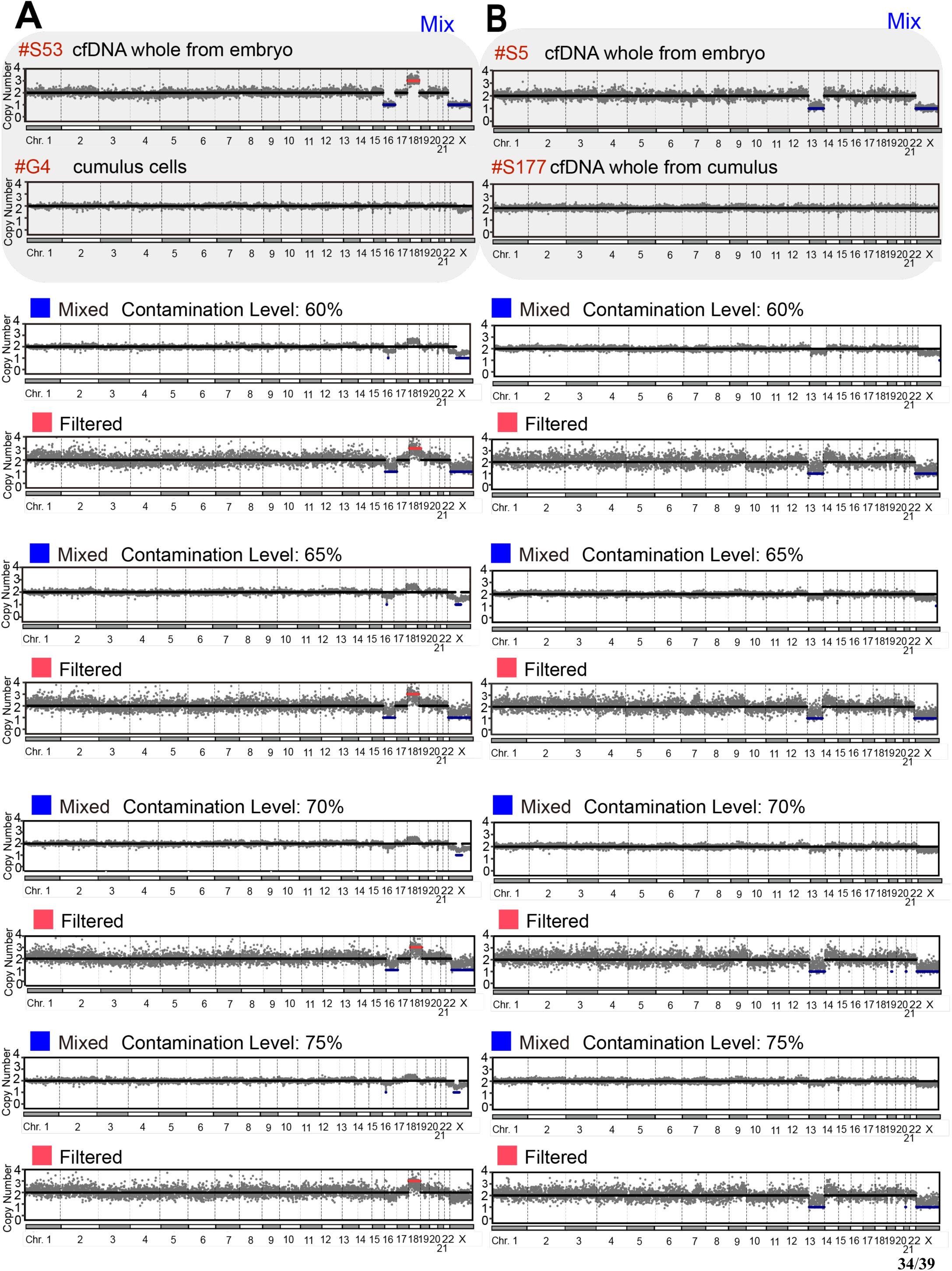
Assessment of Embryonic Chromosome Aneuploidies Reconstruction in Simulated Contaminated SECM Samples. **(A)** Reconstruction of embryonic chromosome ploidy in simulated contaminated samples with varying contamination proportions. Synthetic datasets were generated to mimic scenarios of increasing proportions of cumulus cell contamination (60%, 65%, 70%, and 75%). **(B)** Evaluation of embryonic chromosome aneuploidy reconstruction in simulated contaminated samples with increasing proportions of cumulus cfDNA contamination. Synthetic datasets ranging from 60% to 75% were utilized to simulate contamination scenarios. Blue indicates original mixed samples and red indicates samples that have been filtered after algorithm processing.

Similarly, we began with the other maternal contamination-free aneuploidy SECM sample (S5) and the other SECM sample (S177), which were almost entirely composed of cumulus-derived cfDNA reads (Fig. **4B**, Methods). Again, we introduced cumulus cell reads into the embryonic sample to simulate maternal contamination scenarios at contamination proportions of 60%, 65%, 70%, and 75%. As in the previous simulation, the introduction of contamination led to the obscuration of embryonic chromosome aneuploidy, particularly the seq(13)x1 variation and XY sex, with increasing severity as the contamination proportion increased. Nevertheless, our algorithm successfully reconstructed the seq(13)x1 variation at contamination proportions of 60%, 65%, 70%, and 75%, along with accurate gender identification. Our simulations demonstrated that DECENT could effectively reconstruct embryonic chromosome aneuploidy from contaminated samples up to a certain threshold.

### DECENT reconstructs the embryonic CNV in highly contaminated SECM samples

To test the efficacy of the model in real samples, we applied DECENT to clinical samples with different maternal contamination ratios. We investigated scenarios in which both SECM and TE biopsies were euploid and in which SECM was euploid, but TE biopsy was aneuploid (Fig. **5A**). In the former scenario, despite the culture medium being euploid in the context of high contamination, it does not necessarily imply euploidy in the actual embryo, as maternal contamination could be a contributing factor. Nevertheless, after our algorithmic screening, the results still indicate euploidy, potentially serving as a diagnostic reference.

**Fig 5.**
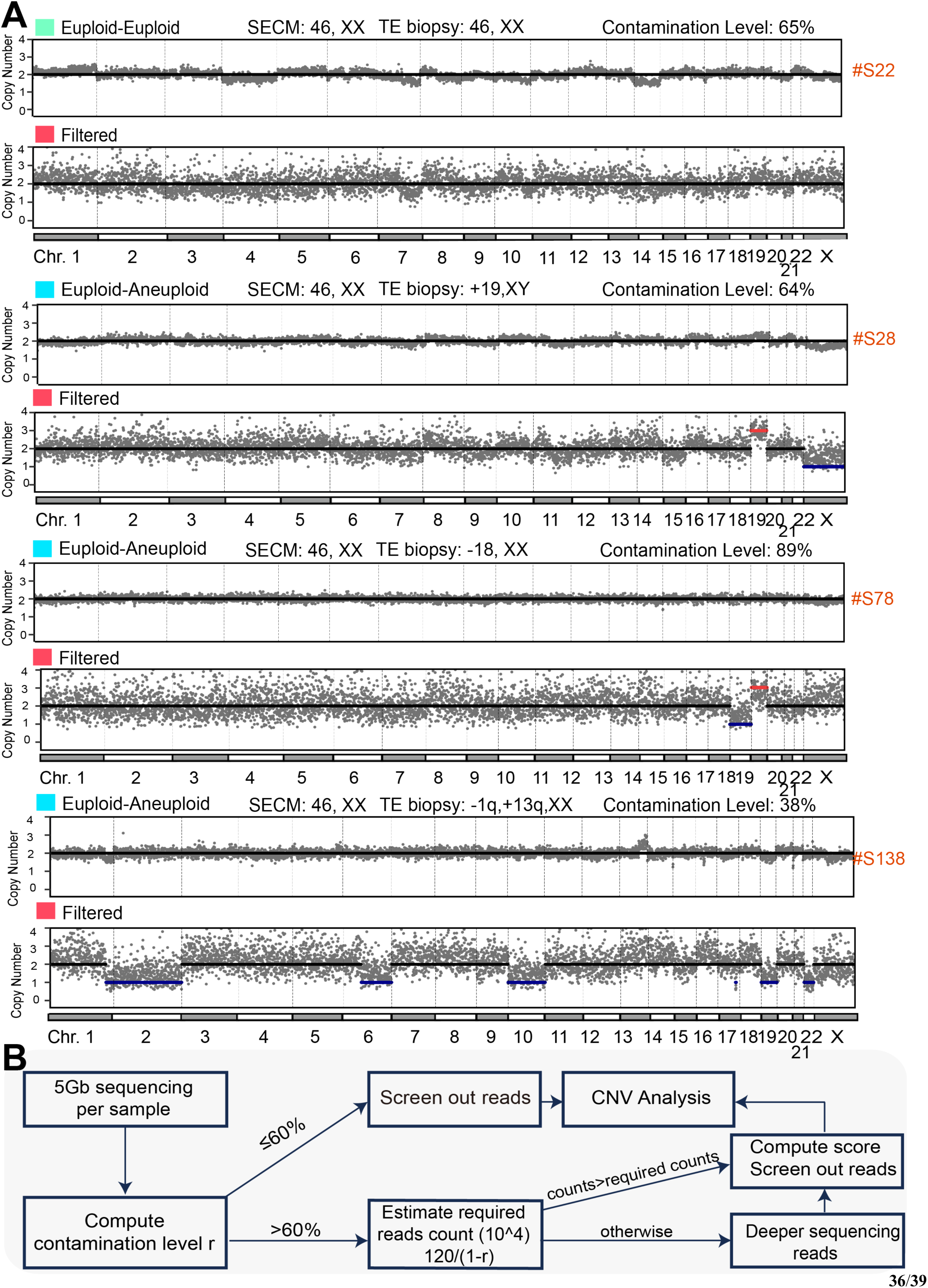
Analysis Results and Clinical Workflow. **(A)** Illustration of our algorithm’s application in reconstructing embryonic CNV from real samples with high maternal contamination proportions (≥60%). Results of DECENT on **S22:** 46, XX; **S28:** seq(19)x3, XY; **S78:** seq(18)x1, (19)x3, XX; **S138:** seq[GRCh37]del(1)(q41q37.3)chr1:g.221575267_243199373del,2x1,del(6)(q12q27)chr6:g.66045183_171115067del, 10x1, 19x1, 22x1, XX. The Light blue block indicates samples with euploid status in SECM and aneuploid status in TE. Cyan block signifies samples that exhibit euploidy in both SECM and TE. Red indicates samples that have been filtered after algorithm processing. **(B)** Workflow depicting the clinical application of our algorithm.

In the latter scenario, false-negative results are represented by maternal contamination, where the presence of chromosome aneuploidy in the embryo is masked by contamination. Through our algorithm, sample S28, with a contamination proportion of 64%, correctly recovered seq(19)x3, XY, which is consistent with the TE biopsy results. Similarly, sample S78, with a contamination proportion of 89%, correctly identified seq(18)x1, aligning with the TE biopsy findings. Notably, in sample S78, in addition to the seq(18)x1 variation, there is also a seq(19)x3 variation, possibly indicating a mosaic phenomenon within this sample. In sample S138, there exist subnormal CNVs, i.e., -1q, +13q in the TE biopsy findings. We find that DECENT also produces the same subnormal CNV. It implies that our method can also handle subnormal CNV cases without only being limited to aneuploidies. We then applied DECENT to all 194 SECM samples with varying degrees of contamination, ranging from low to severe (Supplementary Fig. **S14-S33;** Table **7**).

To facilitate the application of our algorithm in clinical practice, we present a workflow (Fig. **5**B). Since the application of our algorithm leads to a reduction in the number of reads, the decreased read count may impact the final CNV analysis results. Therefore, it is important to ensure that the number of reads after algorithmic screening remains at a certain threshold. To address this, we propose an empirical formula to calculate the number of reads required when the contamination proportion of the sample is *r* (see Methods). Using this empirical formula, we estimate that approximately 3 M reads are needed when the contamination proportion is 60%. We recommend sequencing all samples at least with 5 Gb sequencing per sample. If the contamination proportion exceeds 60%, indicating high contamination, the required number of reads for our algorithm can be estimated using our empirical formula. Subsequently, the sequencing depth of the samples can be determined whether needed to be increased, followed by CNV analysis using our algorithm.

### Filter and attribution analysis reveals potential methylation patterns in embryo development

To further gain biological mechanistic insights from DECENT, we conducted an interpretability analysis of the trained deep learning model. First, we analyzed the features extracted by the first convolutional layer of our neural network, which comprises 100 distinct convolutional kernels (filters). Each kernel independently identifies patterns within the input reads. We visualized the motif features captured by the kernels of our first convolutional layer by calculating their position frequency matrices (PFMs) (Methods). Methylated Cs are now visualized by ‘M’. When ‘M’ and ‘C’ appear simultaneously, it indicates a methylated CpG site in our context. In the analysis, we observed that both methylation and sequence information play significant roles. For instance, in kernels 72, 73, and 74 (Fig. **6A**), the methylation information and sequence information were concurrently influential, indicating that the model may leverage both types of data to make accurate classifications.

**Fig 6.**
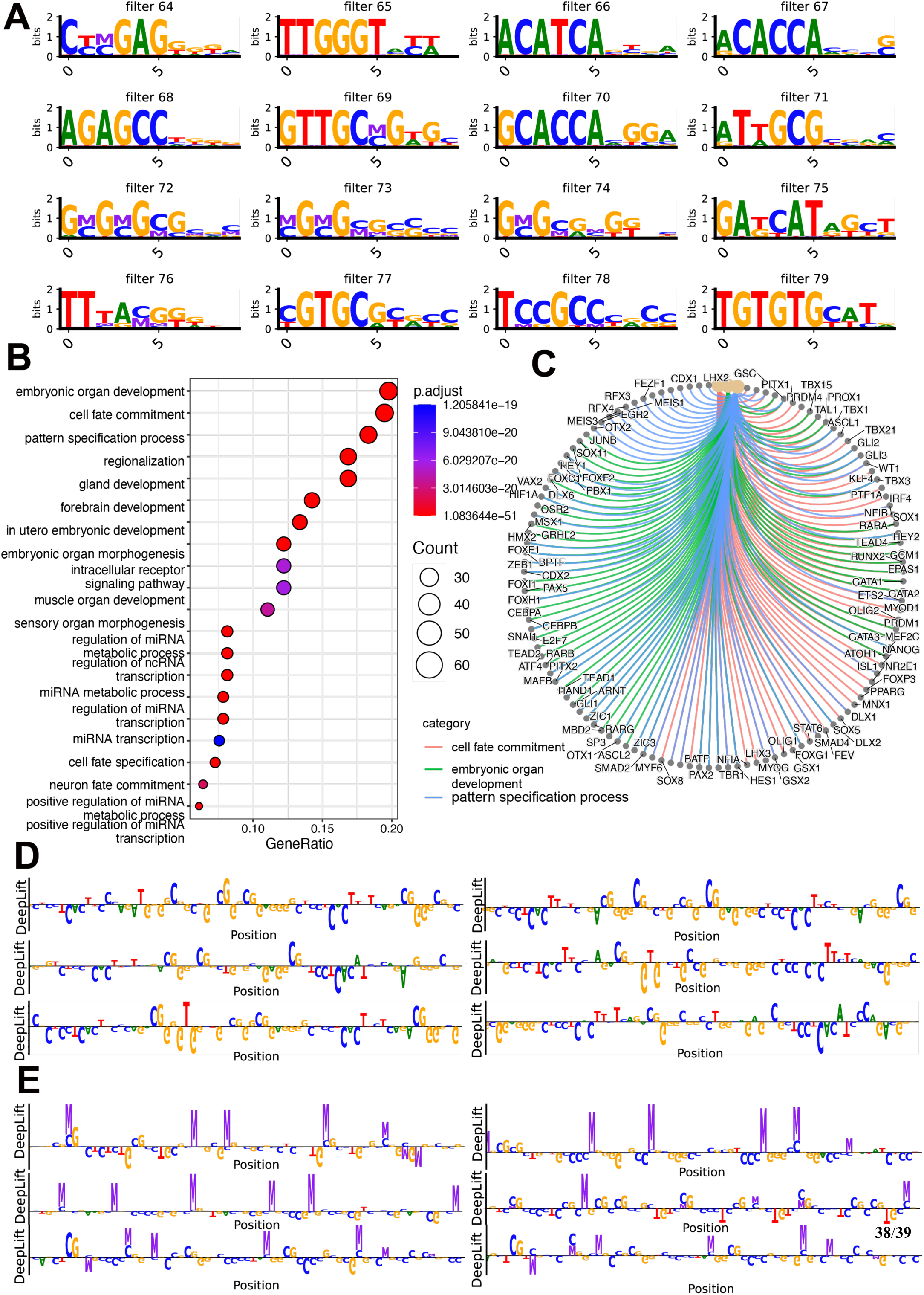
DECENT reveals the underlying patterns and known motifs. **(A)** Visualization of motif features captured by the first convolutional layer kernels (Methylated Cs are visualized by ‘M’. When ‘M’ and ‘C’ appear simultaneously, it indicates a methylated CpG site in our context.). **(B)** Several pathways were revealed by enrichment analysis. **(C)** Visualization of pathways and their associated genes. **(D)** Attribution analysis of reads classified as more embryonic-like by the neural network. **(E)** Attribution analysis of reads classified as more maternal-like by the neural network (the nucleotide at the position of ‘M’ is ‘C,’ indicating a methylated CpG site).

Subsequently, we compared the sequence patterns of these PFMs with known motifs using TOMTOM. We also conducted a gene enrichment analysis using the identified motifs, which uncovered several significantly enriched pathways, particularly those related to embryonic organ development, cell fate commitment, and pattern specification (Fig. **6B**). Since the neural network was trained on data containing both embryonic and maternal origins, the first convolutional layer captures features relevant to both embryonic and maternal processes. With further investigation of the enriched pathways, we observed that the cell fate commitment pathway includes processes such as oocyte fate commitment, the gland development pathway encompasses placental development, as well as the development of structures like the mammary glands and uterus, and the in utero embryonic development pathway involves processes like trophoblast differentiation. These findings indicate that the captured pathways may encompass both embryonic and maternal biological processes. Overall, the filter analysis suggests that the deep learning model may capture biologically meaningful features for classification.

In addition, we conducted attribution (importance) analysis of each nucleotide on reads classified by our model as either resembling maternal or resembling embryonic DNA based on their neural network scores. The analysis (Fig. **6D, E**) revealed that reads with high maternal scores heavily rely on higher methylation information (the nucleotide at the position of ‘M’ is ‘C,’ indicating a methylated CpG site), whereas embryonic reads show lower methylation information. This discrepancy suggests potential differences in methylation levels, implying that the neural network may leverage patterns associated with methylation levels and specific genomic loci for classification. Overall, in addition to its predictive utility in CNV analysis, DECENT is useful for obtaining biological insights into methylation patterns during embryonic development.

## Discussions

In this study, we presented DECENT, a deep learning framework for accurate reconstruction of embryonic CNV in the presence of maternal contamination, a major challenge in noninvasive preimplantation genetic testing. Our method leverages the power of deep learning to effectively capture and model sequence features from both embryonic and maternal cfDNA, enabling contamination level quantification and reliable CNV detection even in scenarios of high maternal contamination.

The performance of DECENT was evaluated using simulated and real datasets. Through simulation analyses, we demonstrated our method can accurately identify embryonic chromosome aneuploidies across varying levels of maternal contamination. Our results showed that our framework could recover embryonic chromosome aneuploidies even in scenarios where the contamination level exceeded 75%. Moreover, validation using real clinical samples further confirmed the effectiveness of our approach, with CNV reconstruction achieved even in samples with high maternal contamination rates. Furthermore, the interpretability analysis conducted on DECENT provided insights into the learning patterns of the neural network. Visualization of convolutional kernel motifs and enrichment analysis of identified genes highlighted the network’s ability to capture potential biologically relevant features associated with embryonic and maternal processes. Additionally, attribution analysis revealed the sequence characteristics utilized by the network for distinguishing between embryonic and maternal reads, further enhancing our understanding of the model’s decision-making process.

Several studies have reported the existence of maternal contamination from non-cumulus cell sources, mainly polar body-source cfDNA, in SECM^12,48,49^. Our earlier findings indicated that polar cell contamination was evident in roughly one-third of SECM cases^22^. However, the incidence of severe polar body contamination in SECM patient samples was minimal, at only 4%. Such contamination can be mitigated by collecting SECMs on day 6 or by postponing the initiation of incubation until day 4^22,50^. Looking ahead, in addition to minimizing polar body contamination during sample collection and processing, we aim to develop new algorithms that incorporate the characteristics of polar bodies, thereby enabling a more effective and comprehensive elimination of polar body-derived contamination.

Meanwhile, it remains challenging to handle uni-parental disomy (UPD) cases with DECENT. We have examined samples with TE biopsy results indicative of UPD and found that the current model cannot accurately recover UPD cases (Supplementary Table **7**). This limitation may not be unique to our algorithm but reflects a broader challenge inherent in next-generation sequencing (NGS) technologies. Without genotyping, NGS has difficulty in accurately detecting UPD^51^. In addition, DECENT could produce some additional small CNV fragments and exhibit limited performance when the contamination ratio is high, and the reads count is low (Supplementary Fig. **S24-S33**). This is due to the relatively low sequencing depth of the current dataset, resulting in fewer filtered reads being available for CNV analysis, which can be mitigated by increasing the sequencing depth and retaining more reads for CNV detection.

As a unified and general method to dissect cfDNA sequences, DECENT could be applied in other tasks beyond niPGT-A, such as noninvasive prenatal testing (NIPT)^52^, non-invasive PGT for monogenic diseases (niPGT-M)^53^, tumor risk prediction, and tissue deconvolution^19^. Pretraining on atlas-scale methylation data^41,54–56^ combined with generative foundation models could also enhance the performance of DECENT. In the future, we will conduct a prospective study and incorporate more additional datasets generated using the same protocol and enhance the sequencing depth for samples with high contamination levels to further validate our algorithm.

Overall, our study contributes to the field of noninvasive preimplantation genetic testing by introducing a new deep learning approach to address the challenge of maternal contamination. By combining an interpretable biological language model, accurate read deconvolution, and CNV reconstructions, our framework has the potential to improve the reliability and effectiveness of non-invasive PGT and potentially facilitate its translation into clinical practice.

## Methods

### Data pre-processing

This work utilized data sourced from the previous study^22^, encompassing a total of 194 PGT-A blastocysts along with their corresponding culture media. The method for detecting DNA methylation in SECM was single-cell whole-genome methylation sequencing. Preprocessing steps involved the removal of sequencing adapters, amplification primers, and low-quality bases from the raw bisulfite sequencing reads. Subsequently, R2 reads exhibiting more than 3 unmethylated CHs, along with their corresponding R1 reads, were discarded. The resulting clean reads were aligned to the human reference genome (hg19) using BS-Seeker2 v2.1.1 (https://github.com/BSSeeker/BSseeker2). Unaligned reads were then remapped to the hg19 genome using the local alignment mode, with the removal of alignments displaying low confidence within microhomologous regions. PCR duplicates were subsequently removed using Picard tools v1.119 (https://broadinstitute.github.io/picard/).

During the training process, six SECM samples with significantly high cumulus contamination levels (>95%) were regarded as cumulus cell source cfDNA, and three samples with zero contamination levels were regarded as embryonic cfDNA based on the DNA methylation signature^22^. Prior to sampling, our dataset consisted of approximately 24,836,198 (∼ 24 million) cumulus reads and 36,846,545 (∼ 36 million) embryonic reads, i.e., in total ∼ 60 million reads. Utilizing the entire dataset for training was not feasible due to practical resource limitations. Specifically, training with tens of millions of reads would require constructing an N × L × 5-dimensional matrix (where N is the reads number and L represents the read length), resulting in a storage requirement exceeding a single 40GB GPU, thereby limiting the feasibility of training both in terms of computational resources and storage capabilities. Additionally, as the training set becomes larger, the training time also increases, making the process even more resource-intensive. So, to address these challenges, we implemented a sampling strategy that balances the need for sufficient training data with the constraints of available computational resources. A total of 15 million reads were randomly sampled from these two categories, with 7.5 million reads each, to form the training dataset. Specifically, 12 million reads (80%) were designated for training, while the remaining 3 million reads (20%) were allocated for validation purposes, ensuring that there was no overlap between these sets.

Then for each sequencing read, we initially trimmed the first 5 base pairs (bp) from the 5′ end to eliminate adapter sequences that could potentially interfere with downstream analysis and model performance. Following this, we uniformly trimmed the 3′ end of all reads to achieve a consistent length of 66 bp. The decision to truncate reads to 66 bp was carefully considered to balance training speed, storage requirements, and model accuracy. The majority of reads have a length of approximately 143 bp. We have conducted experiments comparing the training results using the full length of 143 bp and the truncated length of 66 bp. We found that increasing the read length from 66 bp to 143 bp improved the accuracy by only about 4%. However, this modest gain in accuracy came with a significant increase in training time (4 times increased per epoch), a double increase in storage memory (exceeding one single GPU 40G card memory for training tens of millions of reads), and computational resources required for inference. Therefore, to achieve an optimal balance between performance and computational efficiency, we chose to use reads truncated to 66 bp for our model. We extracted the sequence information and corresponding methylation profile from the reads as inputs for the model.

For our dataset comprising over 15 million reads, training the model for 30 epochs on a single NVIDIA A100 GPU 40G takes approximately 3 hours. The training process is a one-time requirement. Once the model is trained, it can be deployed for inference without the need for additional training or GPU resources. In practical applications, the inference time is often more critical than the initial training time. Applying the trained model to filter a new sample containing over 3 million reads typically takes around 15 minutes on standard computational resources (only use CPU cores).

### Identifying the maternal probability of each read with the deep learning model

To gain a deeper understanding of the influence of sequence information on embryos and cumulus cells, along with the underlying mechanisms, we developed a deep learning model to estimate the probability of sequence reads belonging to maternal origin based on the DISMIR^42^ framework. For the training and validation of this model, we used sequence data obtained from both cumulus- and embryo-derived cfDNA along with their corresponding methylation profiles, which provided additional information for the model to distinguish between reads from the two different sources. We then used one-hot encoding to represent each read in a unified manner. One-hot encoding is a technique used to convert categorical variables into a binary matrix representation, where each category is represented by a vector that has a “1” in the position corresponding to the category and “0” s in all other positions. This encoding involves creating a matrix for each read based on the nucleobase, with an additional encoding for the methylation state. Specifically, each input read is transformed into a matrix of dimensions L × 5, where L is the length of a single sequence read. During the encoding process, methylated Cs at CpG sites were designated in a fifth channel to distinguish them from other Cs. This approach was implemented because CpG sites hold significant biological importance, particularly in humans. They are often located in regions known as CpG islands, which are typically found near gene promoters and play a crucial role in gene regulation. The five channels represent the four nucleobases (A, C, G, T) and the methylation cytosine (mC). For example, if the nucleobase is A, the corresponding one-hot vector is (1,0,0,0,0); for C, the vector is (0,1,0,0,0); for G it is (0,0,1,0,0), for T it is (0,0,0,1,0). For a methylated cytosine (mC), both the ‘C’ and ‘mC’ positions are set to ‘1’, resulting in the vector (0,1,0,0,1). By stacking these nucleotide-specific vectors together for a single read, we form an L × 5-dimensional matrix corresponding to that read. This approach can capture both the nucleotide sequence and the methylation pattern within each read.

The core design of our deep learning approach is conceptually inspired by language models. We aim to leverage analogous methodologies proven effective for language modeling, as sequences and languages share many similarities. We introduce a key modification involving the additional integration of a multi-head attention mechanism. This attention mechanism has proven highly effective in a series of large language models. Specifically, we first use convolutional neural networks (CNNs) to extract sequence features, analogous to “words” in language models. We then integrate an attention mechanism to further capture semantic information within sequences. Subsequently, we employ recurrent LSTM layers alongside CNNs. Finally, a sigmoid activation function is applied to the output, converting it into values within the range of 0 to 1, representing our probability of originating from embryo cells.

Crucial to our method is the integration of the convolution neural network, attention mechanism, and LSTM modules. The CNN captures local information, while the attention mechanism directs interactions at various positions within the sequence, enabling improved extraction of informative details. Furthermore, integration with LSTM enhances our ability to capture and analyze intricate patterns embedded in sequence information.

#### 1) The CNN module

A one-dimensional convolutional layer can extract DNA methylation sequences into latent features by performing the following operations:

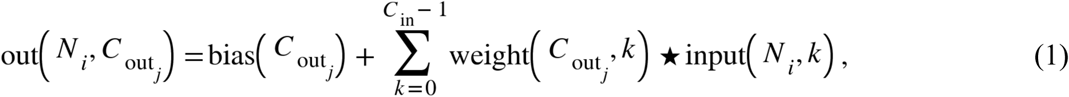

where *N* is the batch size, *C*_in_ and *C*_out_ are the input and output channels, respectively, and ⋆ is the one-dimensional convolution operator. The indices i and j represent the coordinates of the feature map being processed. Specifically, i corresponds to the row (height) position and j corresponds to the column (width) position within the input feature map. The feature integration and extraction can be achieved because the convolutional kernel functions can be considered a sliding window over the sequence, which aggregates local information about the methylation information.

#### 2) The multi-head attention module

The multi-head attention mechanism known as the Transformer^57^ architecture is capable of extracting multifaceted features by simultaneously considering various aspects of the data. In this architecture, input vectors are defined as the query (*Q*), key (*K*), and value (*V*). The mechanism employs a scaled dot-product attention calculation, which is formulated as:

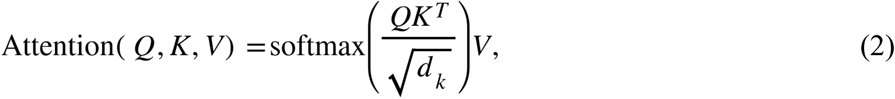

where *Q*, *K* and *V* are the inputs and *d_k_* is the dimension of the key vectors. In the computation of multi-head attention, the input vectors *Q*, *K,* and *V* undergo a series of linear transformations, each corresponding to one of the *h* distinct attention heads (where *h* represents the number of heads). This process involves applying scaled dot-product attention individually to each set of transformed vectors. Subsequently, the resultant vectors from each head are concatenated and subjected to a final linear transformation, resulting in the following formulation:

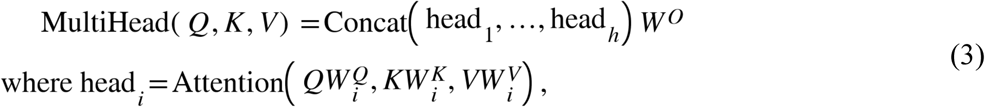

where *W^O^* is the output concatenated weight attention head matrix and 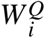, 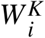 and 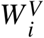 are the weight matrices for *Q, K*, and *V* for the *ith* attention head, respectively.

We have conducted ablation experiments where we removed the attention module from the model (Supplementary Fig. **S13**). The results demonstrated a decline in performance without the attention mechanism, confirming its positive impact on our model’s accuracy.

#### 3) LSTM module

After processing the methylation sequences through convolutional layers and attention mechanism layers, the data is embedded into a feature space. Subsequently, Long Short-Term Memory (LSTM) layers are applied within this embedding space to capture long-term dependencies and temporal patterns in the methylation sequences. This integration of convolutional, attention, and LSTM layers enables our model to effectively learn and model the complex temporal relationships present in the methylation data. The key components of this structure are described as

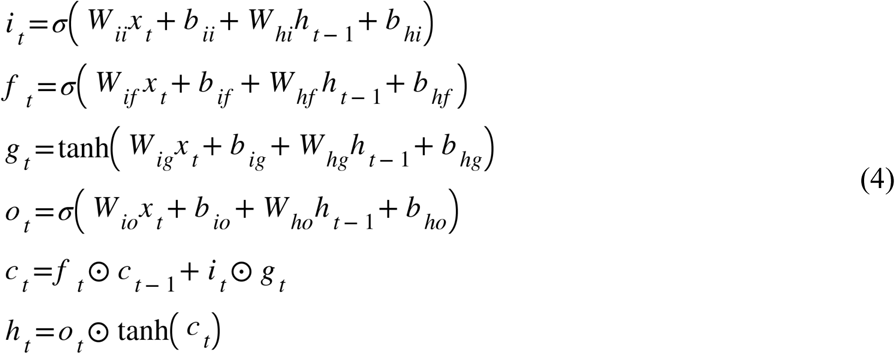

where *W* is the weight matrix. The hidden state at time *t* (*h_t_*) and cell state at time *t* (*c_t_*) are calculated based on the current input *x_t_*, previously hidden state *h_t−_*_1_ (or initial hidden state *h*_0_ at *t* = 0), and gating units, namely, the input gate *i_t_*, forget gate *f_t_*, cell gate *g_t_*, and output gate *o_t_*. The sigmoid function σ modulates the gates, while the Hadamard product applies the gates to control information flow. The integration of LSTM layers enables the modeling of long-range dependencies and patterns in the data sequences. Whereas the preceding CNN and attention layers extract local and contextual features, respectively, the LSTM layers can capture the global sequence structure. LSTM’s gated architecture mitigates vanishing gradients during backpropagation. Moreover, bidirectional LSTM processing incorporates both past and future contexts. Together, the combined power of CNNs, attention, and complementary long short-term memory (LSTM) processing enables robust learning on complex sequential methylation data.

### DMR-based enrichment during training

DMRs are genomic regions exhibiting distinct methylation patterns between different sample groups, and by enriching for reads that overlap these regions, models can leverage these methylation differences to improve their discriminative capabilities.

A total of 769 DMRs were utilized in our study. We sourced a comprehensive set of 27,435 CpG islands (CGIs) from the University of California, Santa Cruz database (genome.ucsc.edu). To mitigate the influence of gender on our analysis, we deliberately excluded CGIs situated on the sex chromosomes. The selection of cumulus-specific CGIs was conducted with precision, adhering to the following criteria: (a) a methylation level exceeding 80% within cumulus cells; (b) methylation levels in other cell types— encompassing sperm, germinal vesicle, MII oocytes, female and male pronuclei, embryos at the 2-cell, 4-cell, and 8-cell stages, morula, inner cell mass (ICM), and trophectoderm (TE)—fell below 20%. These rigorous standards were employed to pinpoint CGIs that exhibit hypermethylation in cumulus cells and hypomethylation in other cell types.

### Reads score range of ambiguous reads exclusion for neural network evaluation

We employed different read score ranges to exclude reads from ambiguous origins whose probability scores were around the central peak of the histogram, ensuring a fair assessment of our neural network. The notation [a,b]\(c,d) is used to describe the inclusion and exclusion of specific score ranges where [a,b] denotes the interval from a to b, inclusive and (c,d) denotes the interval from c to d, exclusive. Therefore, [a,b]\(c,d) means the interval [a,b] excluding the sub-interval (c,d). In practical terms, this includes the range [a, c] and the range [d, b].

We observed that many scores cluster around 0.5, which indicates uncertainty in classification. This clustering arises may because the training dataset is large, and a significant number of reads belong to both the embryo and cumulus categories, making them challenging for the model to classify definitively. So, to evaluate the model’s performance, we implemented a strategy to exclude reads with low confidence scores (around 0.5). By progressively removing reads near the 0.5 threshold, we aim to assess the accuracy of the model in scenarios where it can effectively distinguish between categories ensuring precision. The four thresholds we applied are as follows: 1) No deletion: All reads are included. 2) Deletion of reads with scores in the [0.5, 0.6] range: We remove reads whose prediction scores fall between 0.5 and 0.6. These reads are considered highly uncertain, as a score around 0.5 indicates that the neural network cannot confidently classify the read as belonging to either the embryo or cumulus category. 3) Deletion of reads with scores in the [0.4, 0.6] range: We expand the exclusion range to remove reads with prediction scores between 0.4 and 0.6, retaining reads with higher confidence (scores less than 0.4 or greater than 0.6). 4) Deletion of reads with scores in the [0.3, 0.7] range: We further broaden the exclusion range to remove reads with scores between 0.3 and 0.7, keeping only the reads with the highest confidence (scores below 0.3 or above 0.7). These thresholds were chosen incrementally to assess how removing ambiguously classified reads affects the overall classification accuracy since we are primarily interested in removing this portion of reads.

### Evaluation of model

#### 1) Receiver Operating Characteristic (ROC) Curves and Relevant Metrics

The ROC curve assesses the model’s ability to distinguish between maternal and embryonic reads across all possible threshold values. Here we set the embryonic reads to be the positive class and the maternal reads to be the negative class to ensure better clarity. There are the following components: 1) True Positives (TP): Embryonic Reads correctly identified as embryonic. 2) False Positives (FP): Maternal reads incorrectly classified as embryonic. 3) False Negatives (FN): Maternal reads correctly classified as maternal. 4) True Negatives (FN): Embryonic reads incorrectly identified as maternal. As evaluation metric, the true positive rate and false positive rate can be computed as: 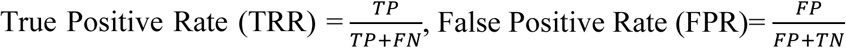. The computation process for ROC is: 1) Threshold Variation: The threshold is systematically varied across the entire [0, 1] range. 2) For each threshold value, reads are classified as embryonic or maternal. 3) Compute TRR and FPR. ROC is depicted as a continuous curve because the points are generated by varying the threshold, thereby illustrating the model’s performance across a range of threshold values. The Area Under the Curve (AUC) is a summary metric representing the model’s overall performance. A higher AUC indicates better discriminatory ability.

#### 2) Precision-Recall (P-R) Curves

The P-R curve illustrates the trade-off between precision (the proportion of true embryonic reads among those predicted as embryonic) and recall (the ability to identify all true embryonic reads) across different thresholds. The precision and recall can be computed as: 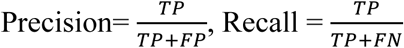 . The computation process for P-R curve is: 1) Threshold Variation: The threshold is systematically varied across the entire [0, 1] range. 2) For each threshold value, reads are classified as embryonic or maternal. 3) Compute Precision and Recall. Similar to the ROC curve, P-R is depicted as a continuous curve because the points are generated by varying the threshold, thereby illustrating the model’s performance across a range of threshold values.

### Computation of contamination levels in the spent embryo culture media

Quantifying contamination levels in SECM is crucial for assessing the reliability of preimplantation genetic testing results. The probability of the deep learning model can be utilized to estimate the extent of cumulus contamination. Such estimations allow for direct comparison with established methodologies, as discussed in^22^, facilitating the evaluation of the efficiency of our approach.

Several estimation techniques are available, among which the maximum likelihood estimation method^41,42^ is a promising approach. Our objective is to derive the contamination proportion within a sample given a set of scores for all *N* reads in the sample. To achieve this, we need to construct the following maximum likelihood function:

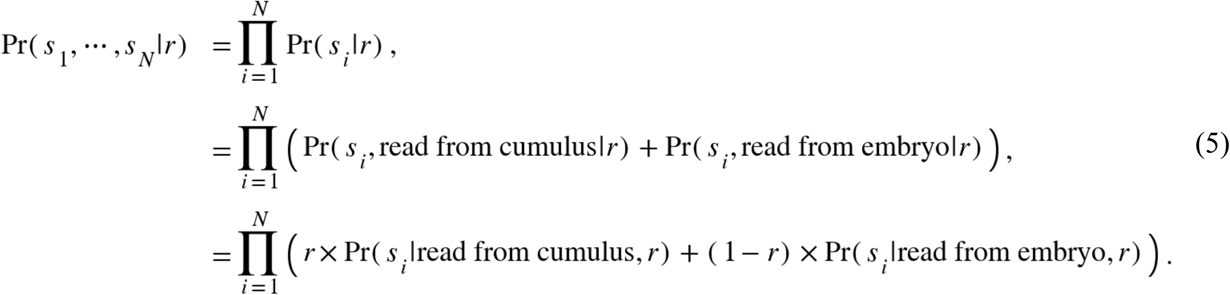

By applying Bayes’ theorem, the posterior distribution of *s_i_* is as follows:

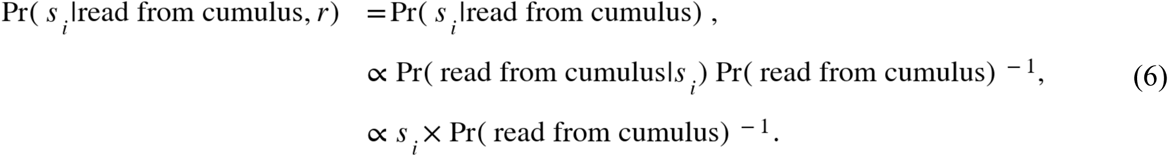

Assuming equal prior probabilities of reads originating from cumulus and embryo, i.e.,

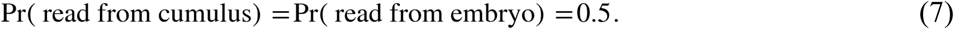

Given this assumption, the likelihood function of *s_i_* can be derived as follows:

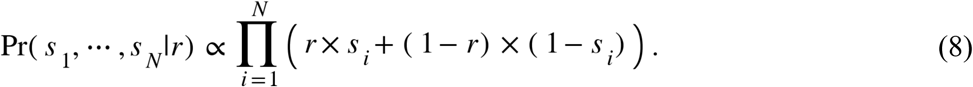

Therefore, the contamination ratio is determined by maximum likelihood estimation:

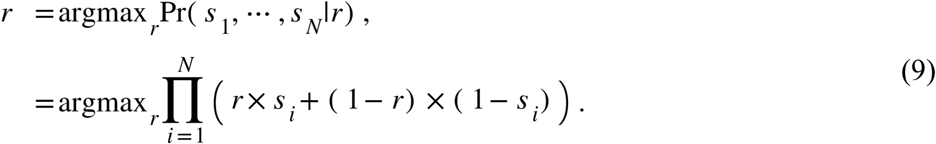

When utilizing Eq. (9) to calculate the contamination ratio, we assume an equal prior probability of reads originating from cumulus and embryo. Hence, in the training set, it is essential to ensure an equal number of cumulus and embryo samples.

### Simulation analysis of read mixture proportions

We conducted a simulation analysis to validate the accuracy of DECENT contamination estimation on synthetic datasets with varying proportions of cumulus and embryonic cell mixtures. The simulation analysis aims to verify that if we take two samples with estimated contamination proportions respectively according to our algorithm, mix them in a predefined ratio, and subsequently estimate the contamination proportion of the mixed sample using the same algorithm, the result is very close to the expected value.

For this purpose, we selected SECM samples with distinct contamination profiles: one sample (S5) with an estimated contamination proportion of 0%, representing reads exclusively derived from embryonic cfDNA, and another sample (S177) estimated almost entirely composed of cumulus-derived cfDNA reads. These samples were mixed in predefined ratios to create a series of synthetic datasets. We performed the mixing of reads randomly and conducted five independent runs of the mixing and estimation process to ensure the robustness of our results and mitigate the effects of randomness. In each run, we introduced Gaussian noise with a fixed mean and a variance uniformly distributed between 0 and 0.1 for each read. DECENT was then used to estimate the proportions of these mixtures based on the read scores. The estimated proportions were compared with the actual mixture ratios to evaluate the performance of DECENT (Fig. **5B**, **5C**). The detailed counts of the mixed reads are provided in Supplementary Table **3**.

### Reconstructing the embryo chromosome CNV

**Simulation analysis:** for the S53 sample (with a contamination proportion of 0, representing reads exclusively derived from embryonic cfDNA) and the G4 sample (composed solely of cumulus cell-derived reads), we first selected approximately 2 million (2M) embryonic-derived reads from the S53 sample. These were subsequently mixed with reads from the G4 sample to simulate varying levels of contamination. Specifically, we introduced 3M, 3.71M, 4.66M, 6M, 8M, and 11.33M G4-derived reads to simulate contamination levels of 60%, 65%, 70%, 75%, 80%, and 85%, respectively. We then applied our algorithm with a contamination threshold of 0.15 for contamination removal.

For the S5 sample (maternal contamination-free aneuploidy SECM sample) and the S177 sample (almost exclusively composed of cumulus-derived cfDNA reads), we selected approximately 2.2M embryonic-derived reads from the S5 sample. These were then mixed with reads from the S177 sample to simulate varying contamination levels. Specifically, we introduced 3.3M, 4.08M, 4.66M, 5.13M, and 6.6M S177- derived reads to simulate contamination levels of 60%, 65%, 70%, 75%, respectively. For higher contamination levels (80% and 85%), due to the limited number of reads available from the S177 sample, we incorporated 8.8M and 12.46M reads from the G4 sample, which is entirely composed of cumulus cell-derived reads. We then applied our contamination removal algorithm with a threshold of 0.15.

**Threshold selection**: the probability score generated by our deep learning network serves as a criterion for filtering embryo reads. As the empirical analysis presented in Figure **2**B, we evaluated the proportion of cumulus contamination across various threshold values using the training dataset. As the threshold decreases, the proportion of reads labeled as maternal (and thus cumulus contamination) also decreases, aligning with our intuitive expectation. At a threshold of 0.15, approximately 90% of the retained reads are correctly identified as embryonic, which meets our accuracy requirements for downstream analysis. Furthermore, to accommodate varying levels of contamination in practical applications, we have provided users with the flexibility to select alternative thresholds based on their specific needs. In our work, we have adjusted our thresholding strategy to better accommodate different levels of contamination. For samples with moderate contamination (10%-60%), we now apply a threshold of 0.2. In cases of low contamination (≤10%), we retain all reads without filtering to facilitate CNV analysis. For high contamination levels (≥60%), we continue to use the stringent threshold of 0.15. Additionally, users retain the option to customize the threshold according to their unique experimental conditions and requirements, allowing for optimized performance tailored to diverse research contexts.

**Inferring CNV**: we used Ginkgo^58^ software for CNV analysis with certain modifications. First, a median length of 500 kb was utilized, with the exclusion of bins that were blacklisted as aberrant. The “500kb” length refers to the size of the variable-length bins used in our analysis. The BED files, derived from the aligned BAM files using bedtools (https://bedtools.readthedocs.io/), served as input files. Genomic GC content bias was mitigated through lowness normalization. For the original samples before filtration, the synthesized BED file derived from randomly extracted normal diploid blastocyst reads was utilized as the reference. This reference set serves as a baseline for comparing the test samples and identifying variations, ensuring that the observed differences are not due to technical biases or noise and is utilized in Ginkgo analysis. For filtered samples, the reference sample utilized in the original study was filtered by our algorithm with a threshold of 0.15. The resulting samples were converted to BED files to serve as references for the filtered samples. Utilizing cytoband data obtained from the UCSC Genome Browser, we calculate the bands and absolute positions of the regions where CNVs occur. When CNVs encompass an entire chromosome, this is characterized as aneuploidy. Conversely, when CNVs are confined to specific segments of a chromosome, they are referred to as sub-chromosomal CNVs.

Subsequently, after applying our algorithmic filtering, the number of reads was reduced, which made the final CNV visualizations appear noisier. To address this and enhance the clarity of the visualizations, we applied a variance reduction technique to the CNV data points post-Ginkgo processing. This step was solely intended to improve the visual presentation and did not involve diminishing any signals. To reiterate, the CNV analysis performed by Ginkgo comprises the following key steps: (1) binning reads into genomic regions; (2) quality control through read coverage and uniformity analysis; (3) removal of outliers, normalization, GC correction, and bin smoothing; and (4) segmentation of bins and determination of copy number states.

**Estimation of required read counts for CNV reconstruction**: when the contamination proportion calculated by the DECENT algorithm is denoted as r, the estimated total counts of the required reads are approximately Counts=120/(1-r) (10k). The number 120 (10k) is derived from empirical testing conducted during the development of our algorithm. Specifically, we aimed to identify the optimal number of reads necessary for reliable CNV analysis across different contamination levels. Through our investigations, we observed that when the contamination ratio is 0%, our method retains approximately 10% of the original reads using a threshold of 0.15 (Supplementary Table **6**). Given that CNV analysis typically requires a minimum of 100,000 reads to achieve reliable results, we set the baseline at 120,000 reads (or 120 counts, in units of 10,000) to provide a buffer for variability in read quality and distribution. So given the contamination ratio r, the formula 120 / (1 - r) was derived to scale the number of retained reads proportionally based on the estimated level of contamination. Specifically, when *r* is 60%, the required number of DECENT reads is approximately 3 million. However, we suggest that deeper sequencing is preferred if it is possible.

### Filter and attribution analysis of the deep learning model

To gain deeper insight into the knowledge acquired by the model and to elucidate the underlying mechanisms governing the sequence reads, we embarked on interpretability analysis of the trained deep learning models. This analysis aims to unravel the complex relationships and decision-making processes inherent within the model’s architecture. Here, we use EUGENe^59^ (version 0.1.2), a Python toolkit, to interpret our sequence-based deep learning models.

First, when identifying specific features recognized by each filter within the first convolutional layer, we use approximately 100,000 typical maternal reads (high score) and 100,000 typical embryonic reads (low score) from the SECM samples, totaling 200,000 reads for analysis. This involved calculating the activation values associated with the weight matrices, thereby offering insights into the filter-specific feature detection and representation capabilities of the model. Subsequently, the extracted feature motifs were compared to certain motifs from the human genome using TOMTOM^60,61^ (version 5.5.5). We employed a sliding window approach, examining all possible 10-base pair (10bp) subsequences across the reads to ensure comprehensive motif identification. This comparison enabled us to identify genes enriched with these motifs. For these enriched genes, we conducted gene enrichment analysis using clusterProfiler^62^ (version 3.19) to uncover the pathways in which these genes are involved.

Subsequently, to quantitatively ascertain the contribution of individual nucleotides within the input sequence to the predictive outcome of the model, we conducted an attribution analysis. In this analysis, we employed the DeepLIFT^63^ method implemented in EUGENe, which is designed to provide a detailed understanding of feature influence in deep neural network predictions. This approach is effective for elucidating which sequence features play a pivotal role in the neural network’s discriminative capabilities.

### Statistical analysis

Python (version 3.10) and R software were used for statistical analysis. A Student’s t-test was used in **Fig. 3A, 3C**. Statistically significant comparisons were shown, with p < 0.05 as significant. The sample size (n=194) for statistical analysis was described in the legend of the corresponding figure.

## Acknowledgments

We sincerely appreciate the support from grants from the National Key R&D Program of China (2023YFC2705600, 2023YFC2705602 to Y.C.), the National Natural Science Foundation of China (82301889 to Y.C. and 12288101, 8206100646, T2321001 to P.Z.), Key Clinical Projects of Peking University Third Hospital (BYSYZD2022029 to Y.C.), Young Elite Scientists Sponsorship Program by CAST (2023QNRC001 to Y.C.), Beijing Natural Science Foundation (7232203 to Y.C.), Start-up Grants of Peking University (7101303365 to P.Z.) and Peking University Medicine Sailing Program for Young Scholars’ Scientific & Technological Innovation (BMU2023YFJHPY001 to Y.C.). We are also thankful for the support from the High-performance Computing Platform of Peking University and the Computing Platform of the Center for Life Science for data analysis. We are grateful for the helpful suggestions from Professor Tiejun Li on this work.

## Author contributions

Y.C. and P.Z. conceived the project; Z.Z. and P.Z. designed the algorithm; Z.Z. implemented the algorithm and performed the computations; all authors analyzed the data and interpreted the results; Z.Z. drafted the initial manuscript with input from all authors; all authors revised and approved the manuscript.

Y.C. and P.Z. supervised the project.

