## Supplemental information for "Genetic deconvolution of embryonic and maternal cell-free DNA in spent culture medium of human preimplantation embryos through deep learning"

### Supplementary Information

### A Supplementary Figures

#### A.1 Supplementary Figure S1: Evaluation of DECENT on Validation Dataset

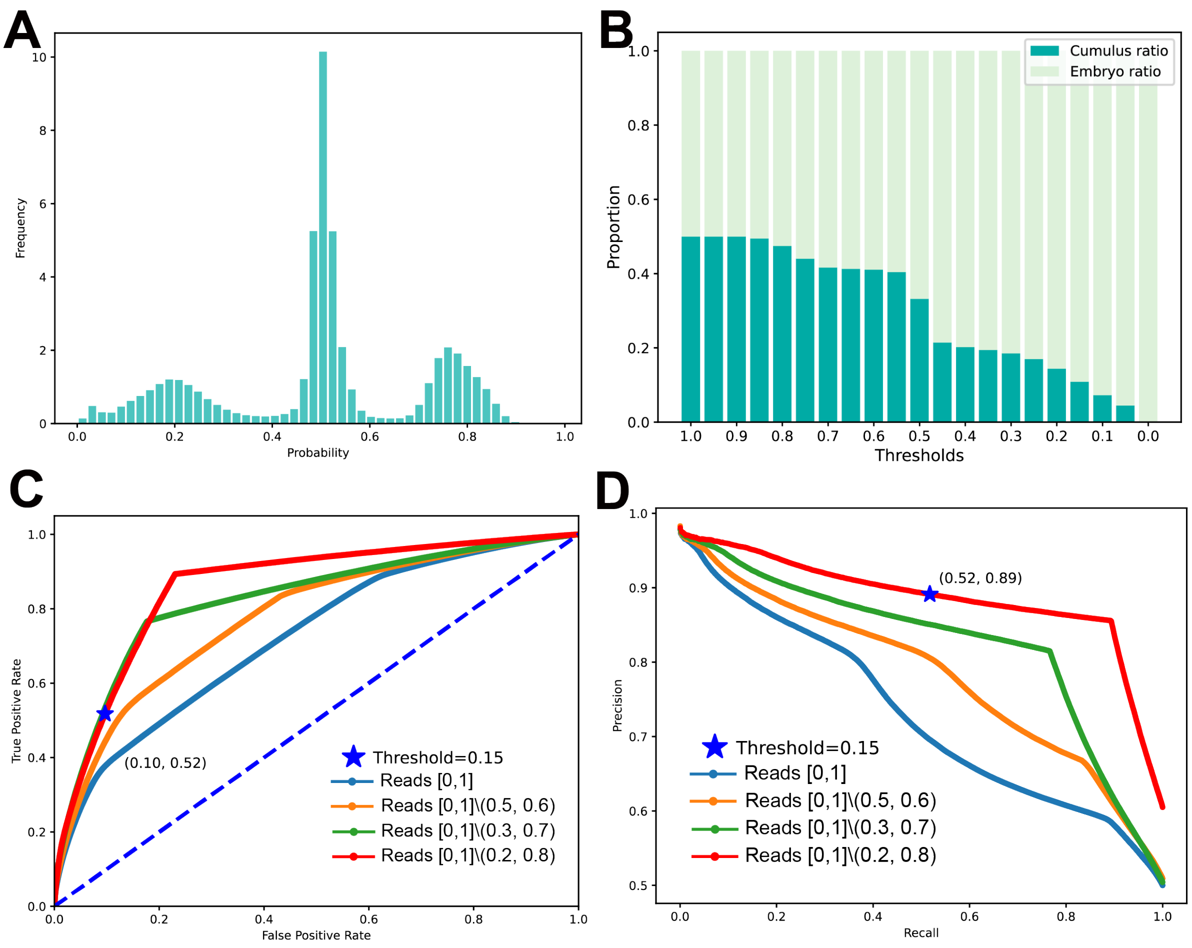

**Figure S1.** Evaluation of DECENT on the validation dataset. **(A)** Distribution of scores generated by the neural network model on the validation dataset, showing three main modes concentrated around 0.2, 0.5, and 0.8, respectively. **(B)** Variation in identified cumulus contamination proportion with different threshold values on the training dataset. A decrease in the proportion of cumulus contamination is observed as the threshold decreases. **(C)** Receiver operating characteristic (ROC) curves comparing the performance of the deep learning model under different threshold ranges for read exclusion. **(D)** Precision-recall (P-R) curves demonstrating the trade-off between precision and recall for the deep learning model under different threshold ranges for read exclusion.

#### A.2 Supplementary Figure S2: Evaluation of Embryonic CNV Reconstruction in Simulated SECM Samples with Contamination Levels Exceeding 75%

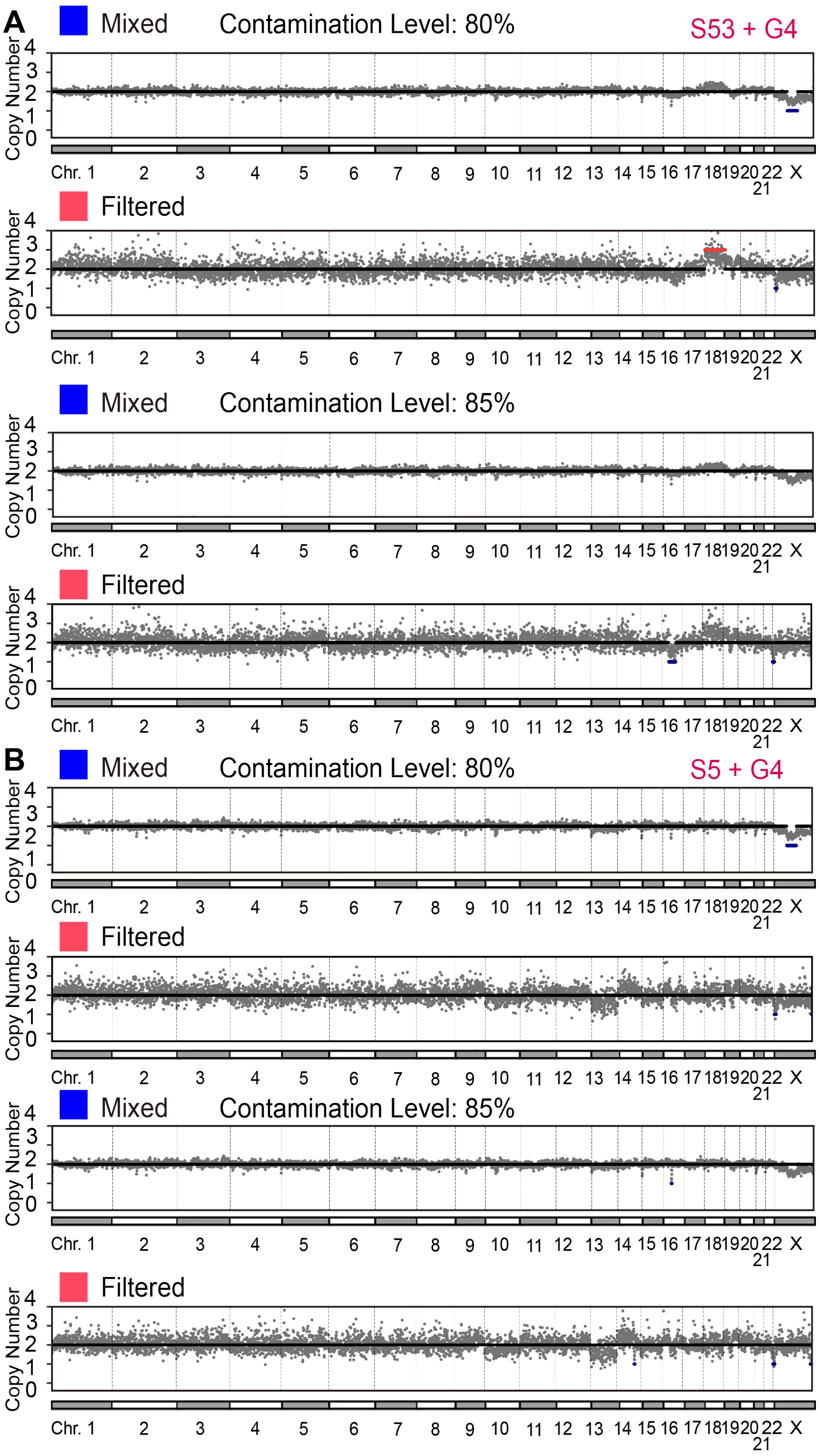

**Figure S2.** Evaluation of embryonic CNV reconstruction in simulated SECM samples with contamination levels exceeding 75%. **(A)** S53 and G4, **(B)** S5 and G4.

##### A.3 Supplementary Figure S3: Influence of Reads Count on CNV Analysis Results

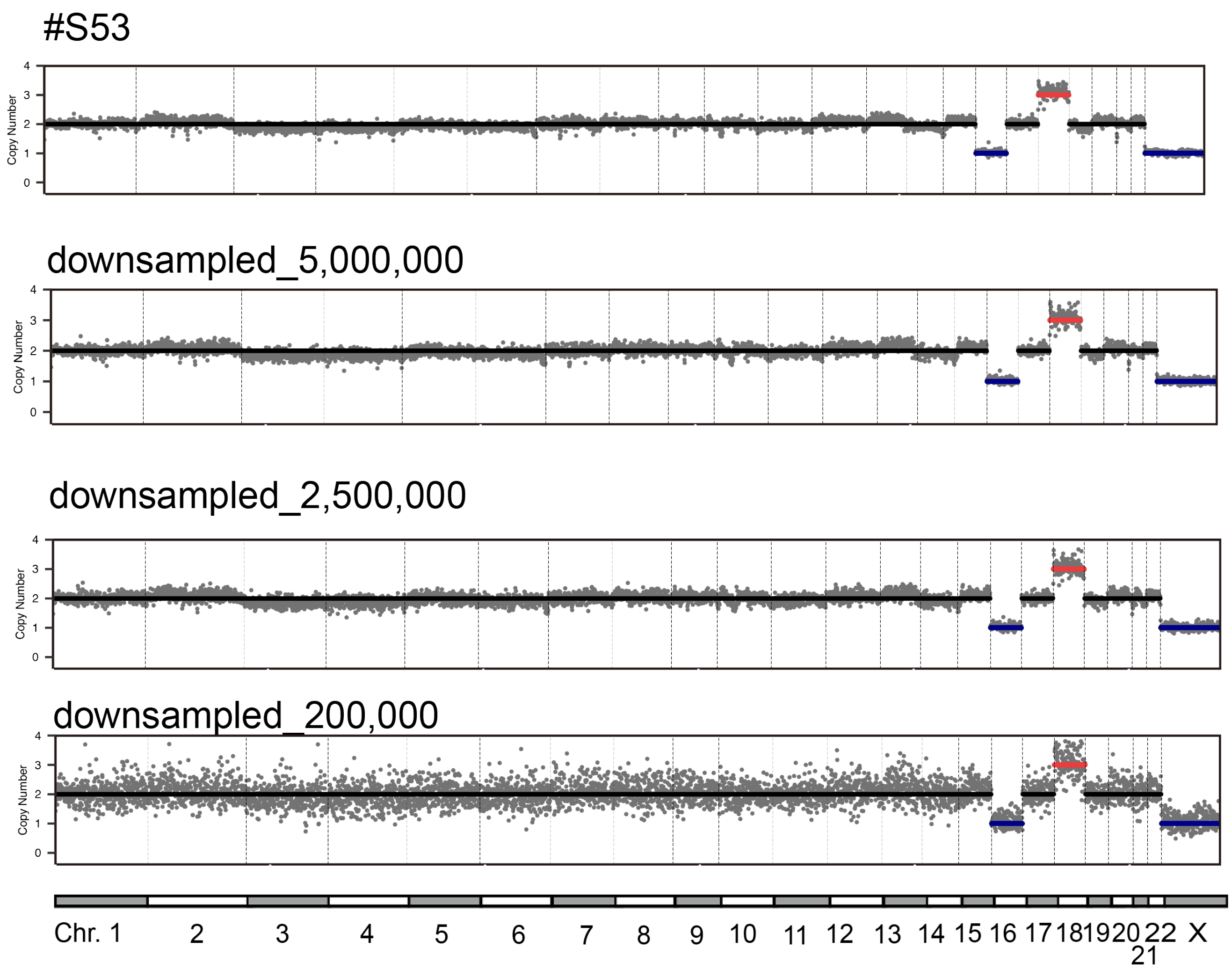

**Figure S3.** Influence of reads count on CNV analysis results. We selected sample S53 which CNV have -16, +18 profile. Then the reads were downsampled to certain counts respectively, to investigate the effect of read count on CNV analysis results.

##### A.4 Supplementary Figure S4: Filter Analysis of DECENT (I)

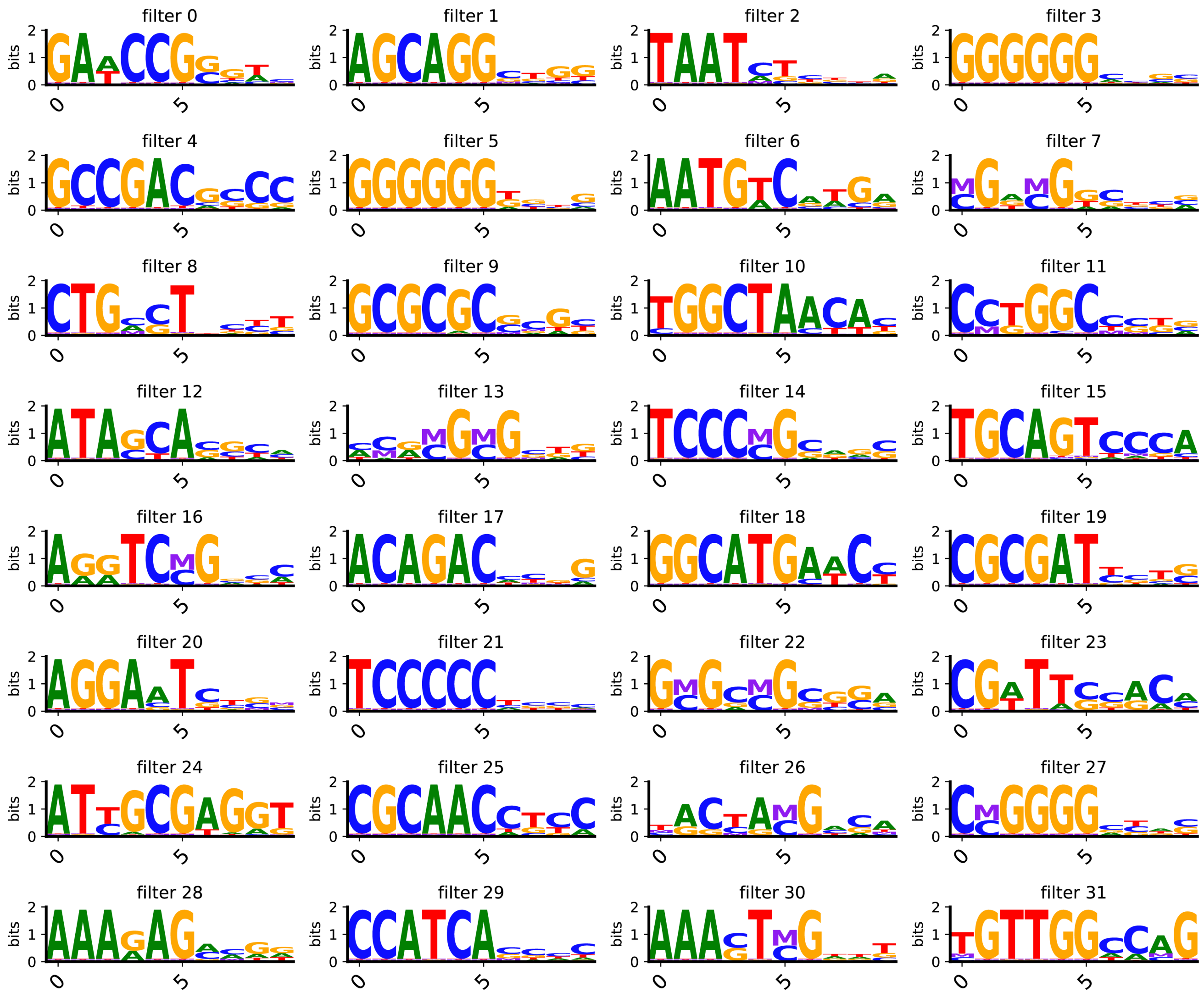

**Figure S4.** Filter analysis of DECENT (I). Visualization of motif features captured by the first convolutional layer kernels.

##### A.5 Supplementary Figure S5: Filter Analysis of DECENT (II)

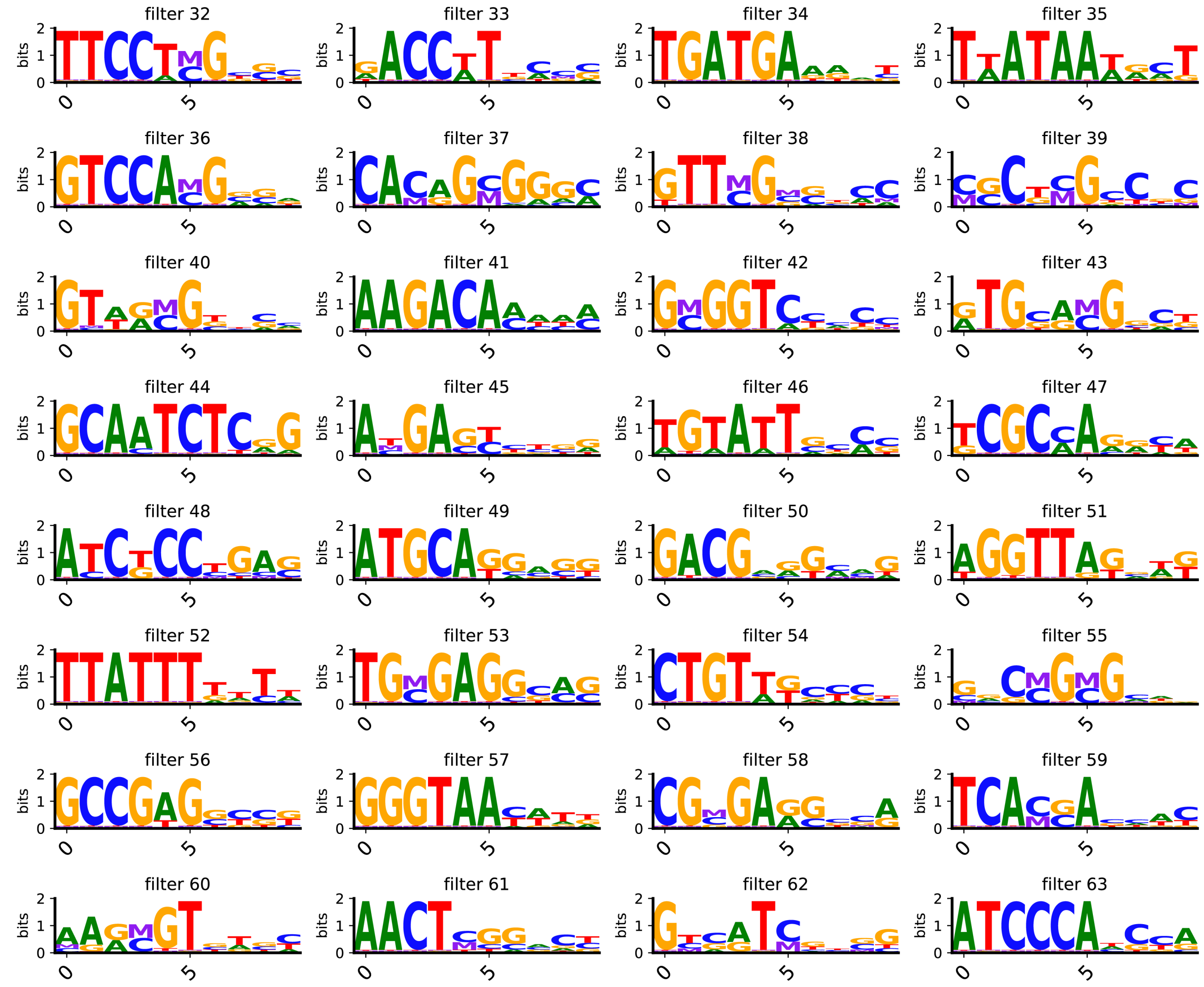

**Figure S5.** Filter analysis of DECENT (II). Visualization of motif features captured by the first convolutional layer kernels.

##### A.6 Supplementary Figure S6: Filter Analysis of DECENT (III)

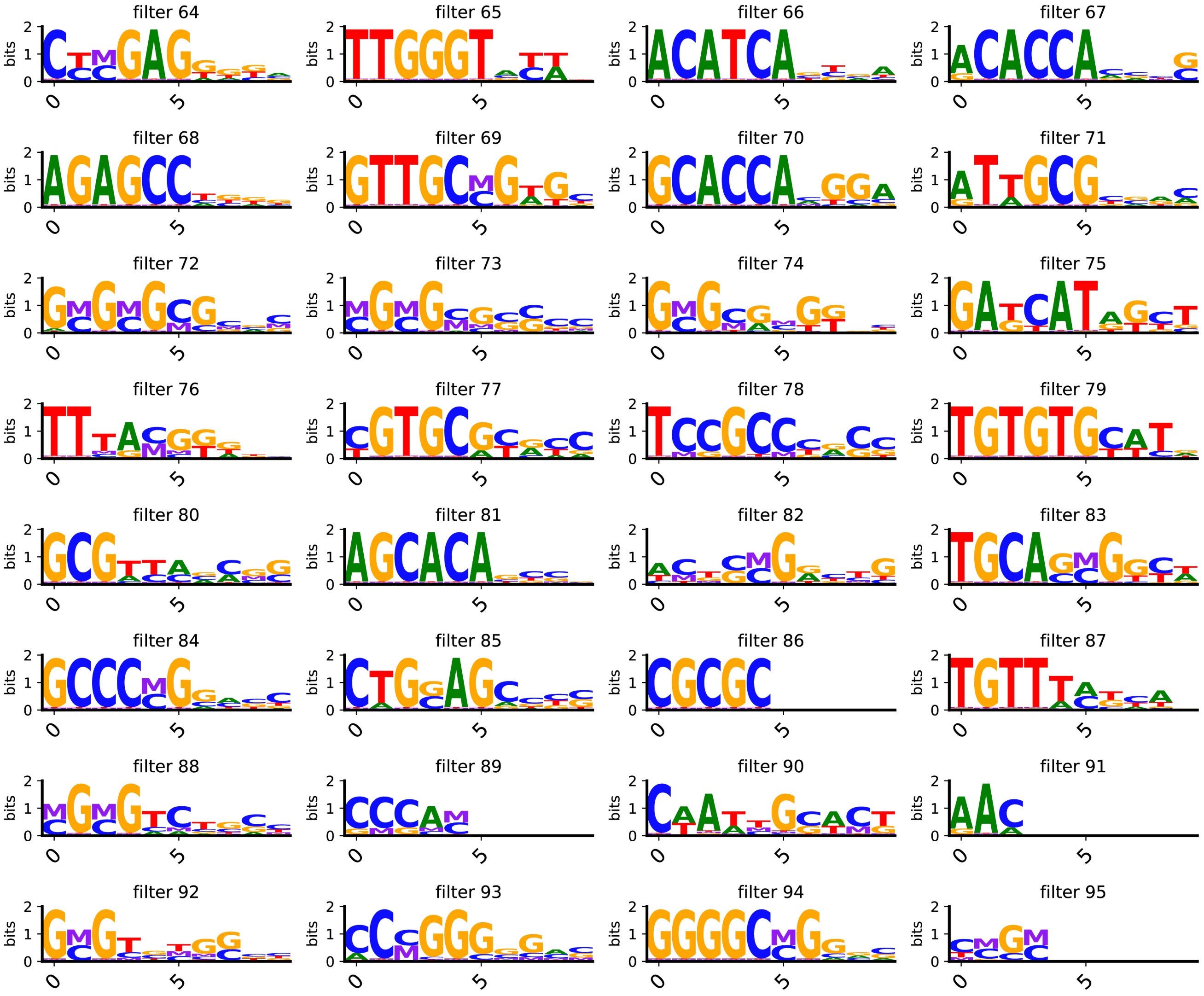

**Figure S6.** Filter analysis of DECENT (III). Visualization of motif features captured by the first convolutional layer kernels.

##### A.7 Supplementary Figure S7: Attribution Analysis of DECENT on Embryonic Reads

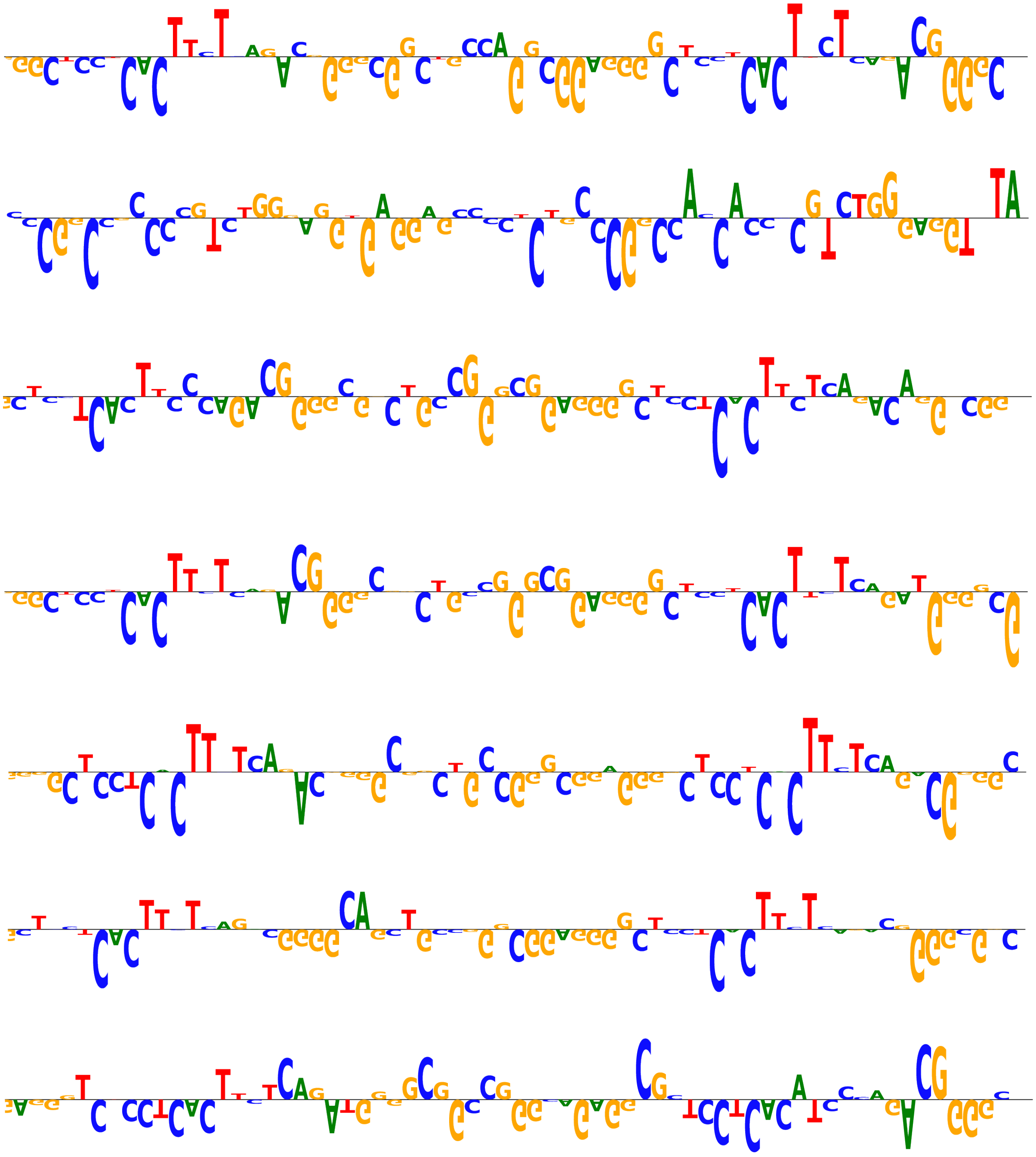

**Figure S7.** Attribution analysis of DECENT on reads classified as more embryonic-like by the neural network.

##### A.8 Supplementary Figure S8: Attribution Analysis of DECENT on Cumulus-Liked Reads

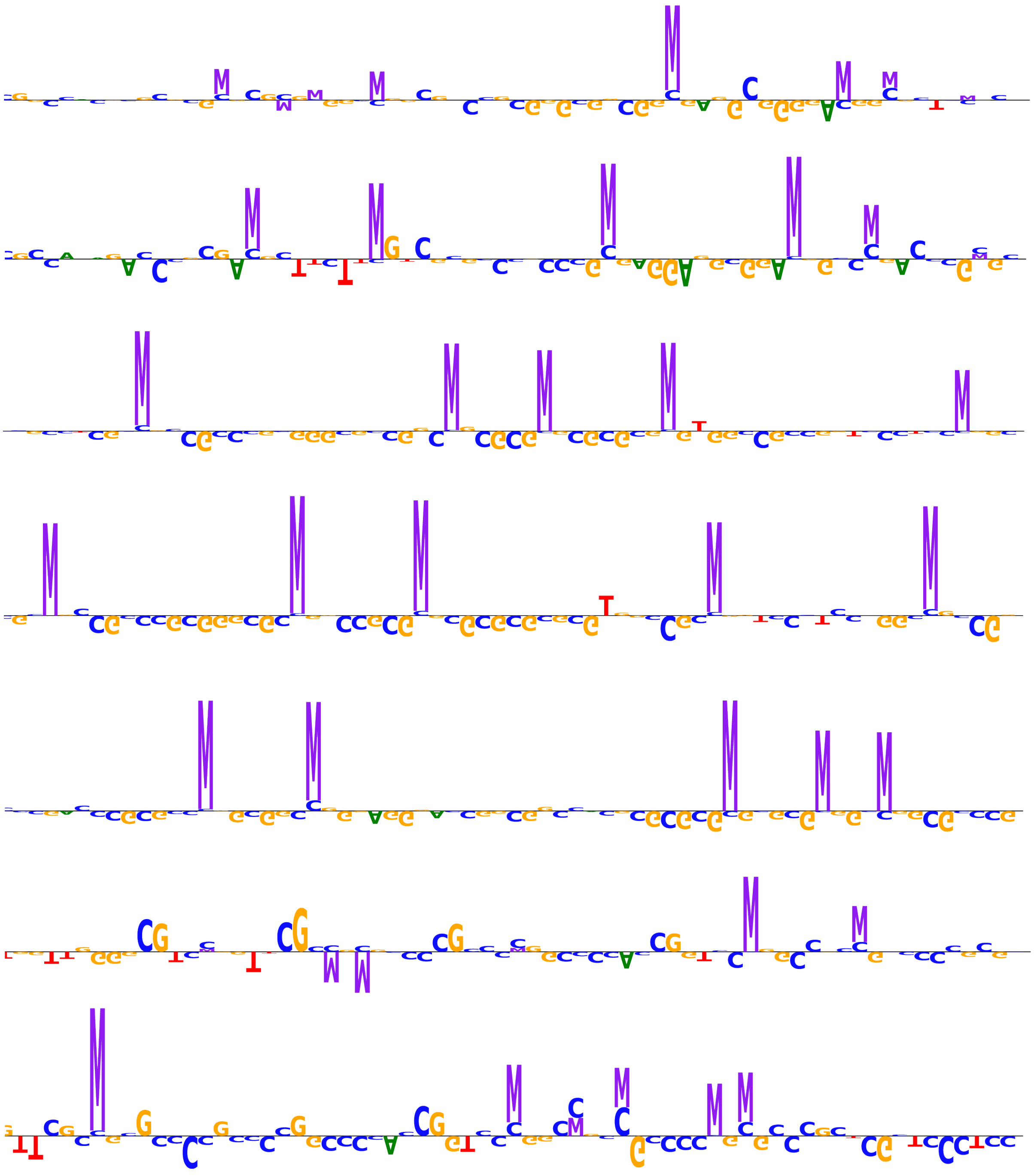

**Figure S8.** Attribution analysis of DECENT on reads classified as more maternal-like by the neural network.

##### A.9 Supplementary Figure S9: Distribution of high-scoring maternal reads and low-scoring embryonic reads in PCA space across different neural network layers.

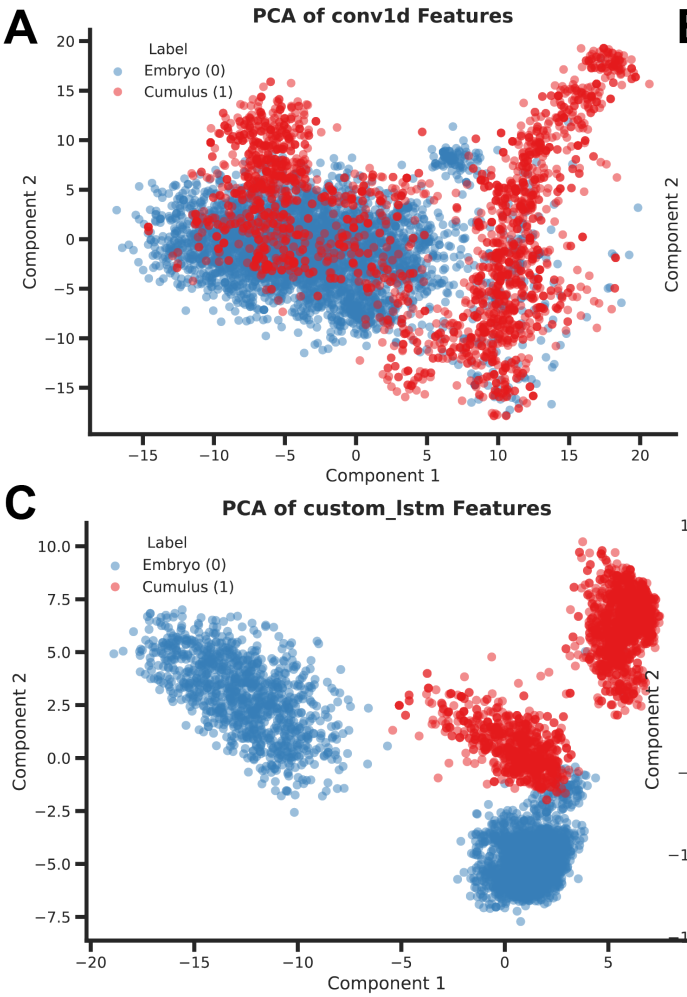

**Figure S9.** Distribution of high-scoring maternal reads and low-scoring embryonic reads in the principal component space across different neural network layers. (A) The first convolution layer. (B) The attention layer. (C) The LSTM layer. (D) The second convolution layer.

##### A.10 Supplementary Figure S10: Attention Weight Distribution of DECENT on Embryonic Reads

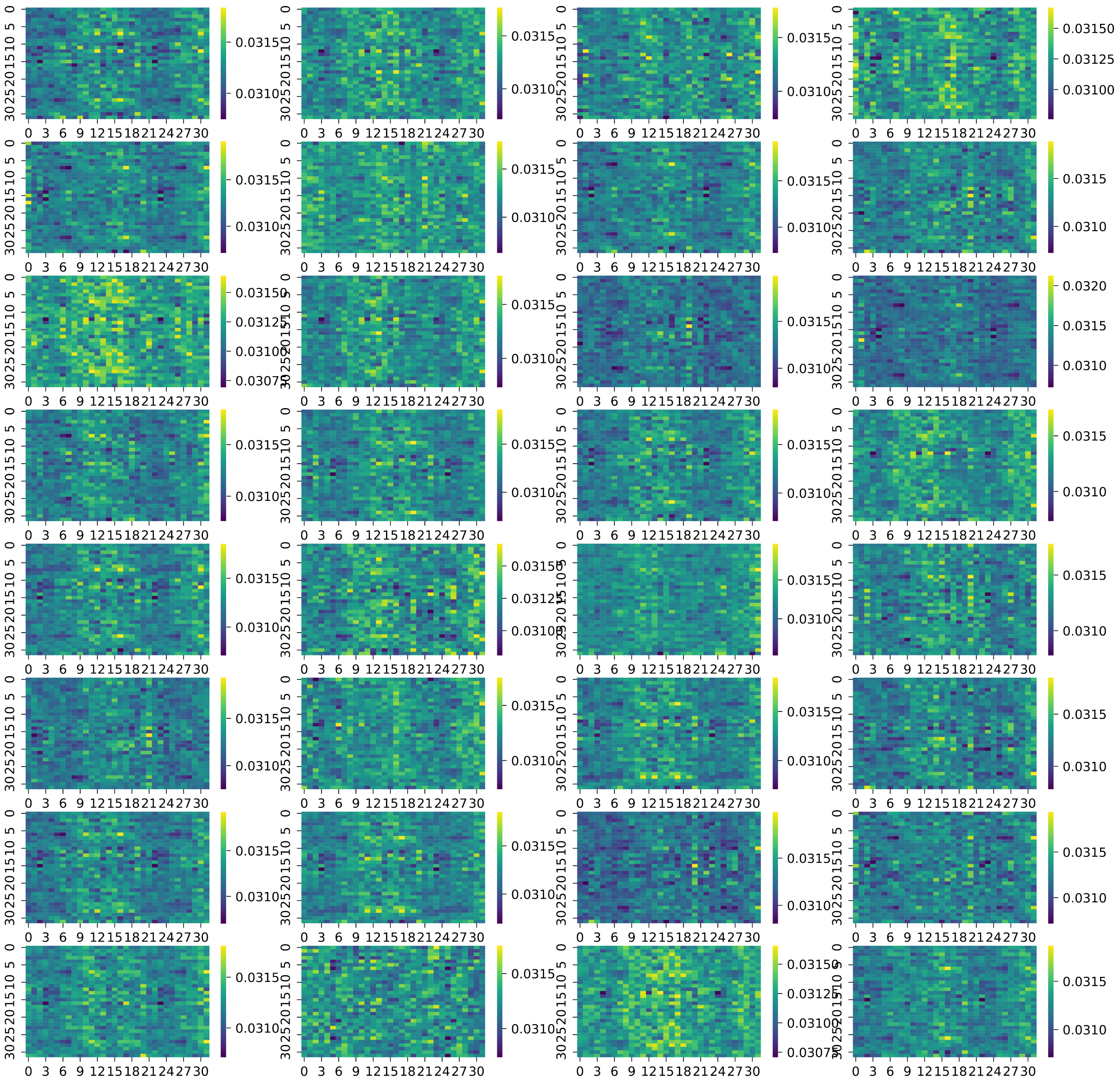

**Figure S10.** Attention weight distribution of DECENT on embryonic reads. Visualization of average head attention weight matrices captured by the multi-head attention module.

##### A.11 Supplementary Figure S11: Attention Weight Distribution of DECENT on Cumulus-Like Reads

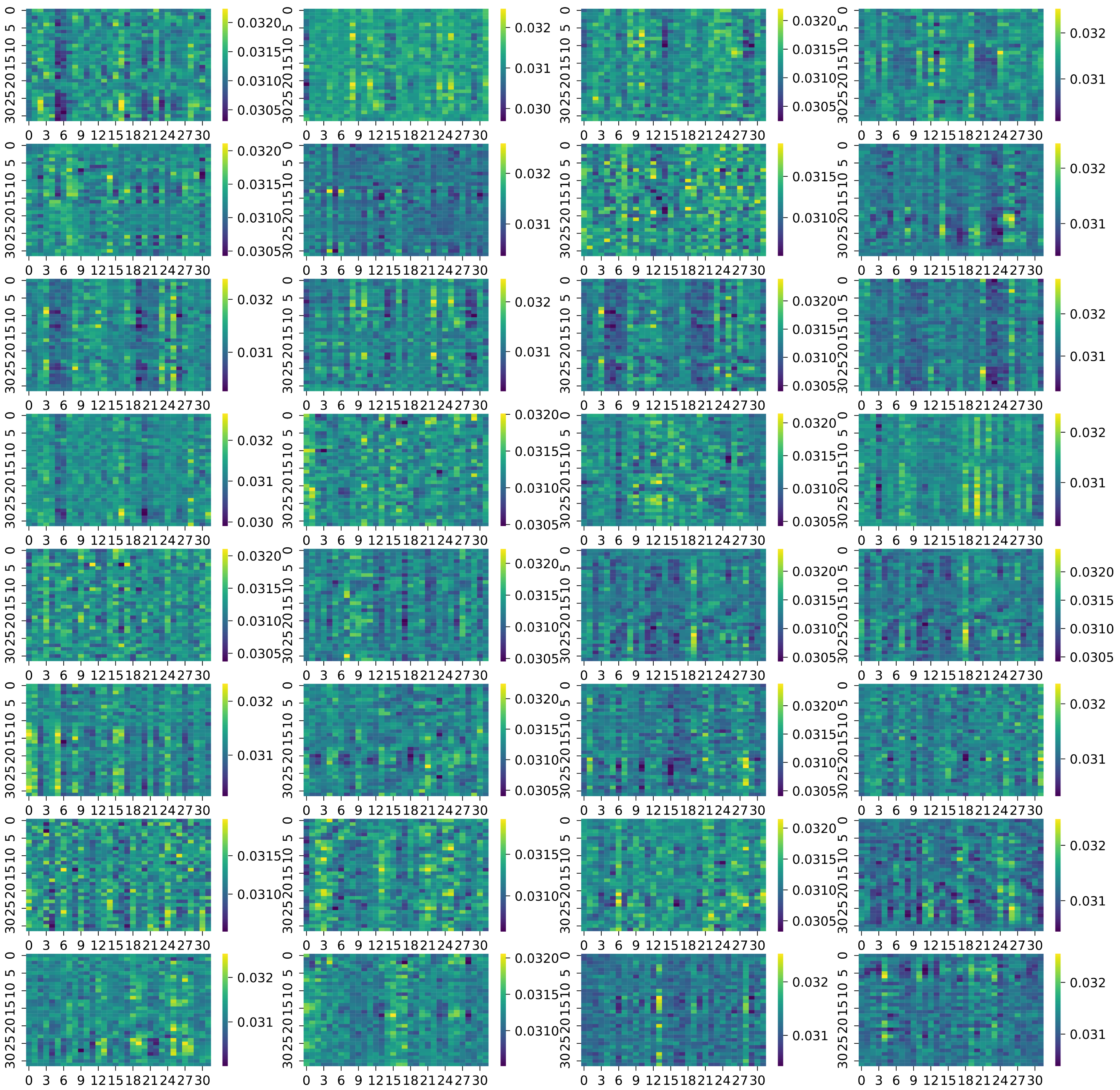

**Figure S11.** Attention weight distribution of DECENT on cumulus-like reads. Visualization of average head attention weight matrices captured by the multi-head attention module.

##### A.12 Supplementary Figure S12: Effect of DMR Enrichment on Model Classification Performance.

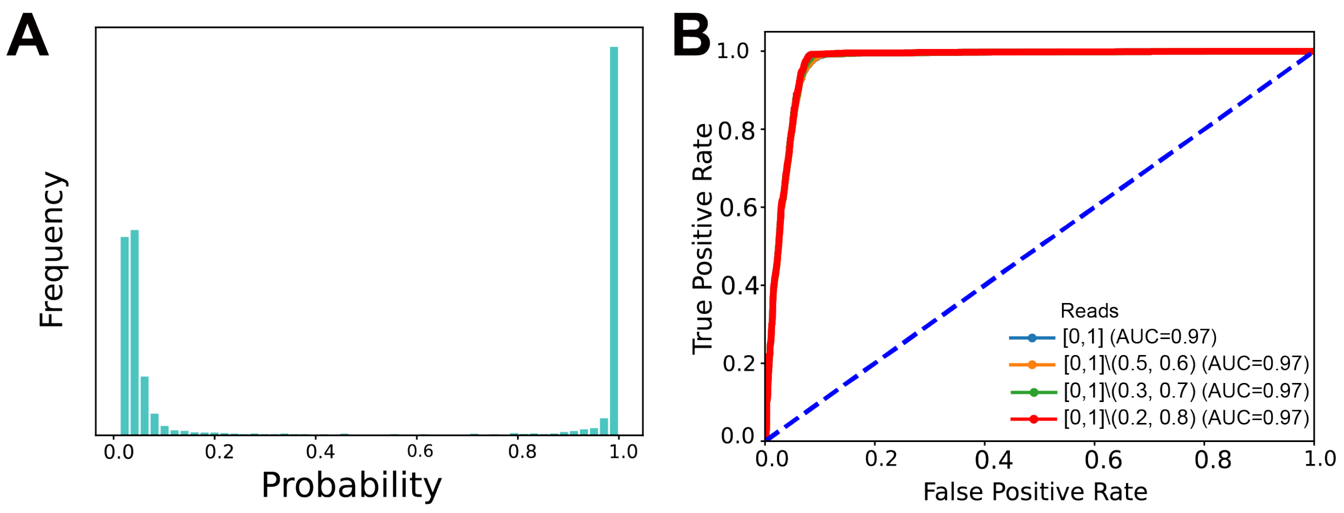

**Figure S12.** Effect of Differentially Methylated Regions (DMR) Enrichment on Model Classification Performance.

##### A.13 Supplementary Figure S13: Training loss and validation loss versus epochs with or without attention.

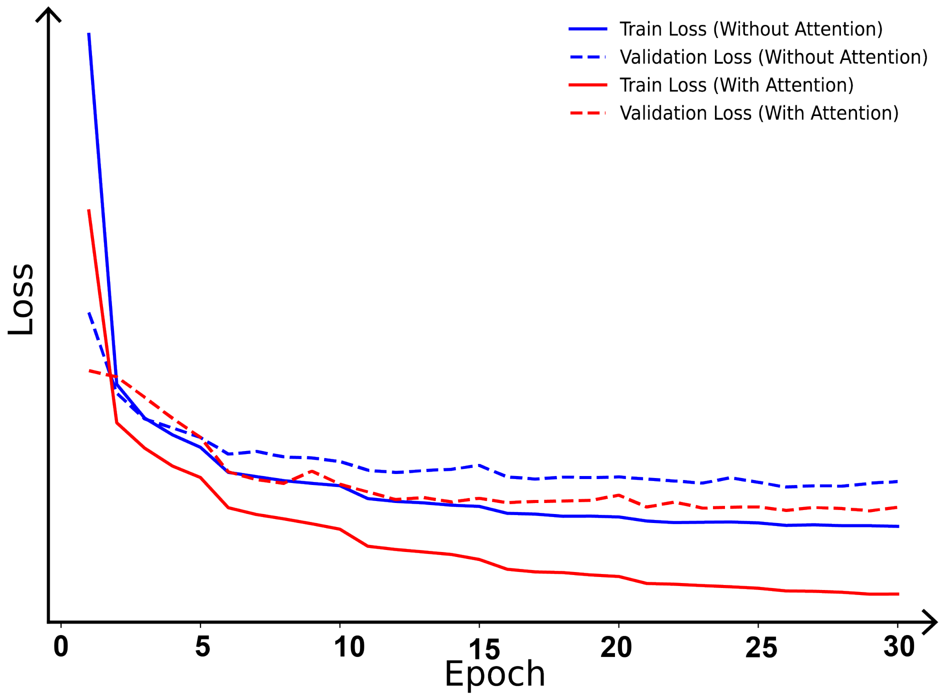

**Figure S13.** Training loss and validation loss versus epochs with or without attention.

#### A.14 Supplementary Figure S14: Analysis Results of DECENT on Moderate Contaminated SECM samples (I)

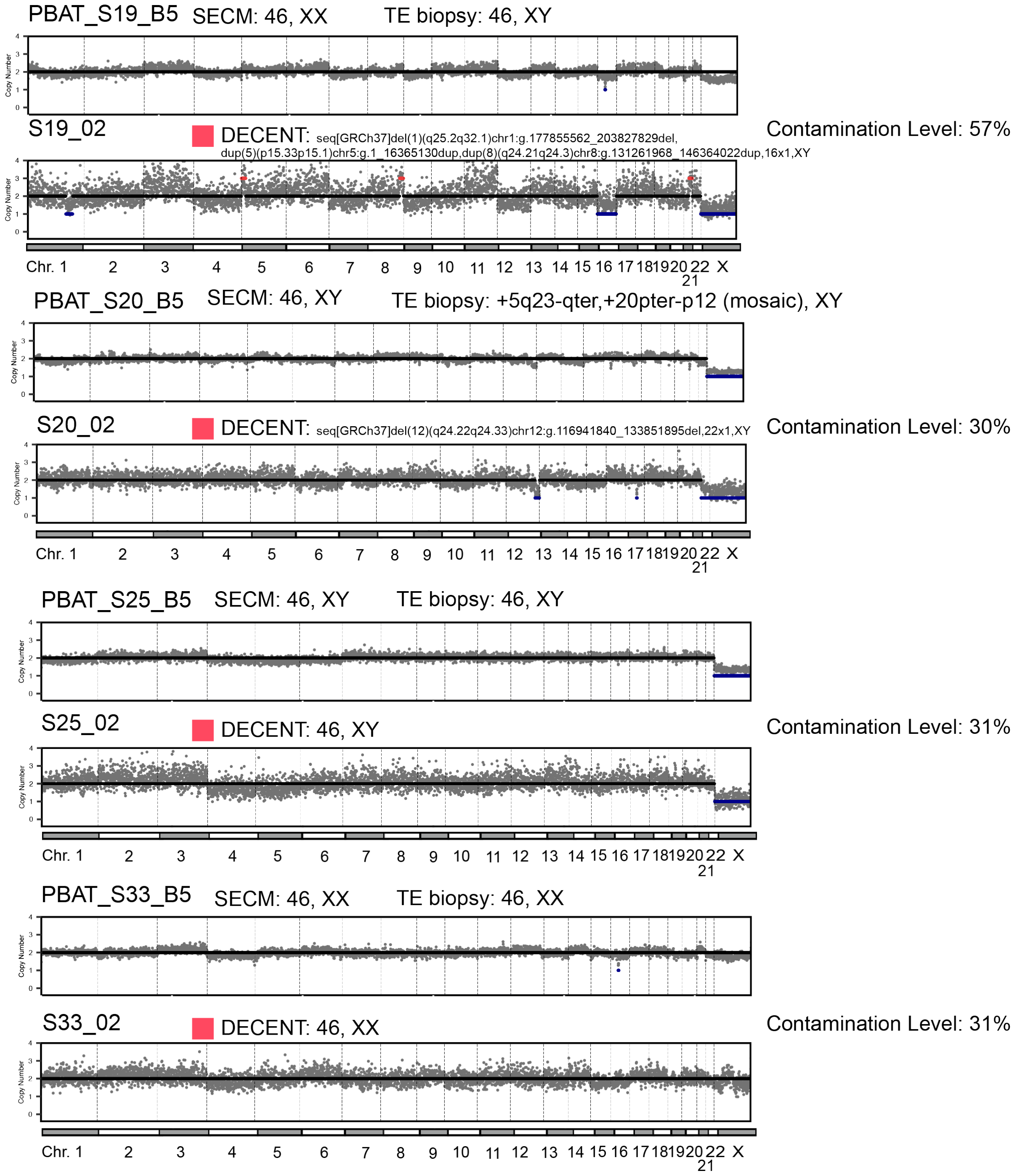

**Figure S14.** Analysis results of DECENT on moderate contaminated SECM samples (I). Each pair of plots represents a sample set, with the first plot displaying the chromosome copy number results of the original sample and the second plot showing the chromosome copy number results after applying DECENT filtering with a threshold of 0.2. We show the original SECM, TE and the processed DECENT results.

#### A.15 Supplementary Figure S15: Analysis Results of DECENT on Moderate Contaminated SECM samples (II)

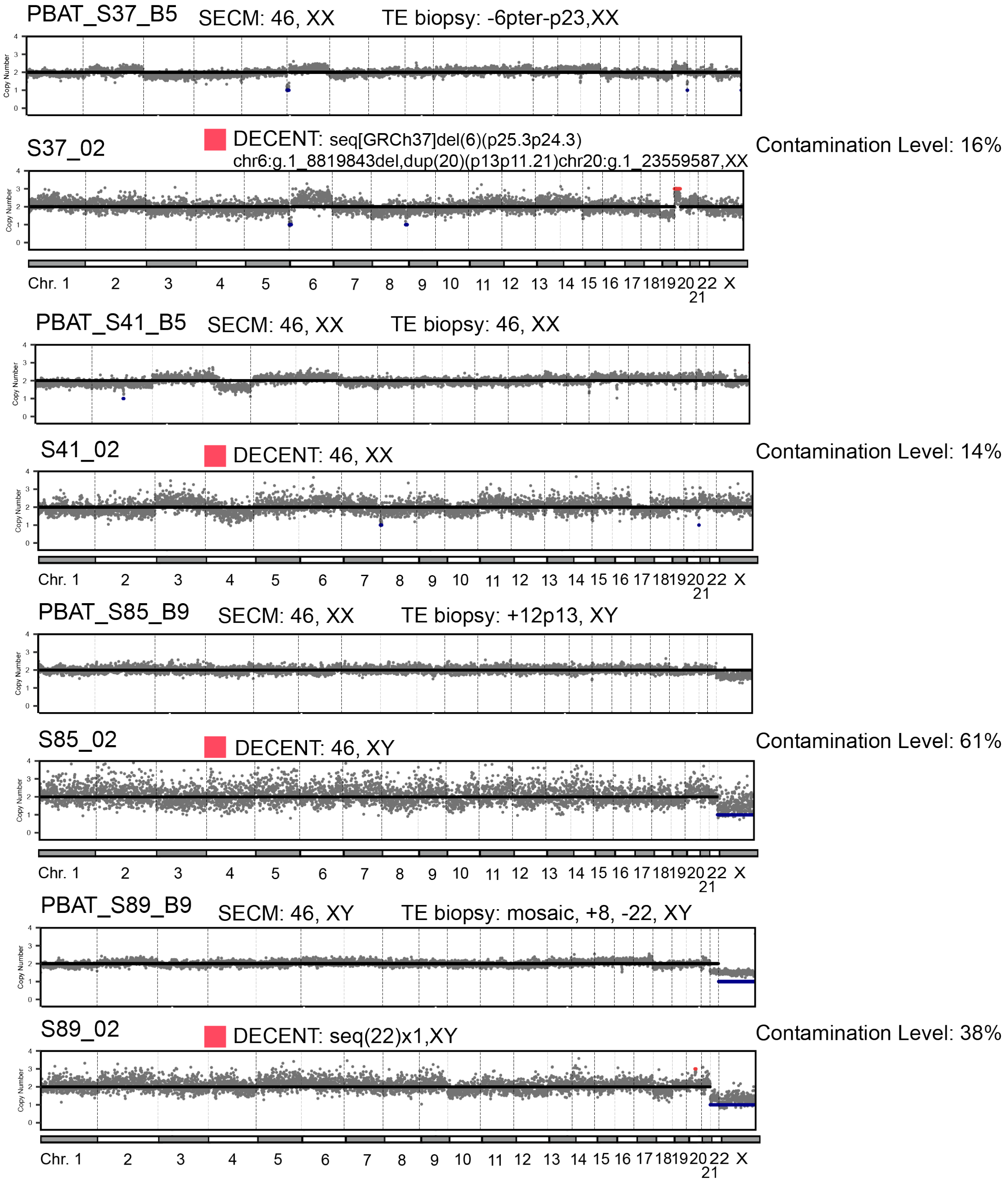

**Figure S15.** Analysis results of DECENT on moderate contaminated SECM samples (II). Each pair of plots represents a sample set, with the first plot displaying the chromosome copy number results of the original sample and the second plot showing the chromosome copy number results after applying DECENT filtering with a threshold of 0.2. We show the original SECM, TE and the processed DECENT results.

#### A.16 Supplementary Figure S16: Analysis Results of DECENT on Moderate Contaminated SECM samples (III)

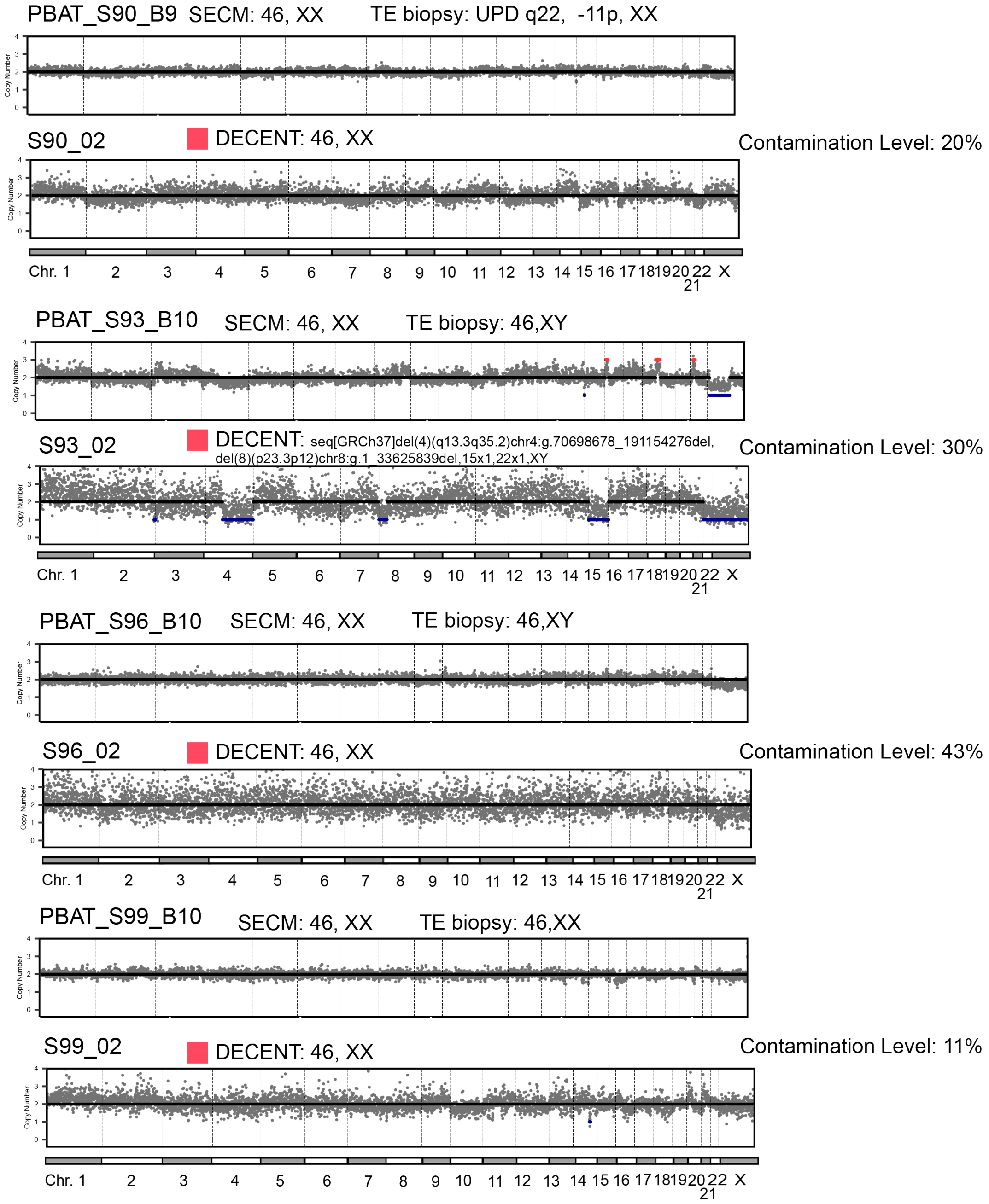

**Figure S16.** Analysis results of DECENT on moderate contaminated SECM samples (III). Each pair of plots represents a sample set, with the first plot displaying the chromosome copy number results of the original sample and the second plot showing the chromosome copy number results after applying DECENT filtering with a threshold of 0.2. We show the original SECM, TE and the processed DECENT results.

#### A.17 Supplementary Figure S17: Analysis Results of DECENT on Moderate Contaminated SECM samples (IV)

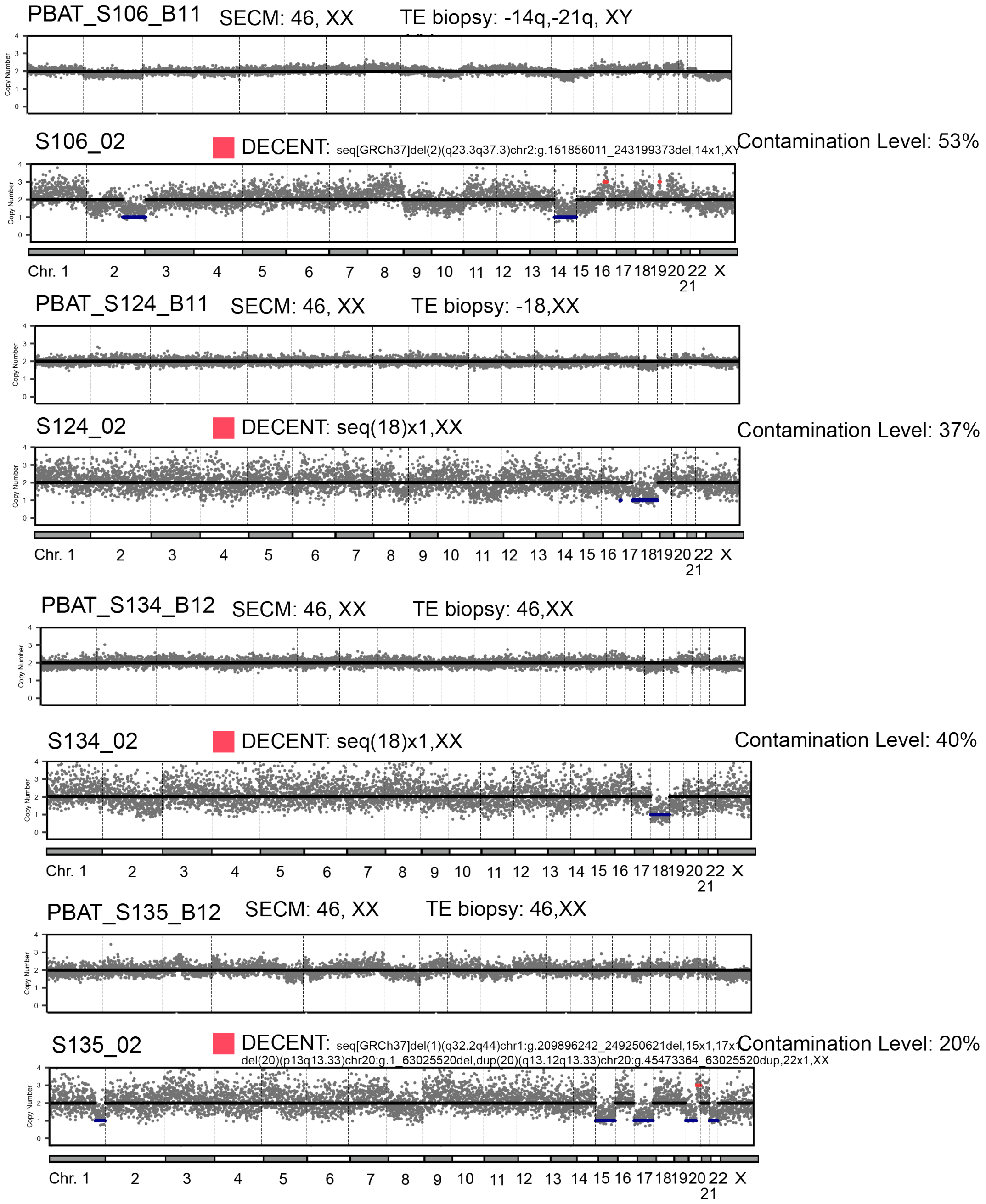

**Figure S17.** Analysis results of DECENT on moderate contaminated SECM samples (IV). Each pair of plots represents a sample set, with the first plot displaying the chromosome copy number results of the original sample and the second plot showing the chromosome copy number results after applying DECENT filtering with a threshold of 0.2. We show the original SECM, TE and the processed DECENT results.

##### A.18 Supplementary Figure S18: Analysis Results of DECENT on Moderate Contaminated SECM samples (V)

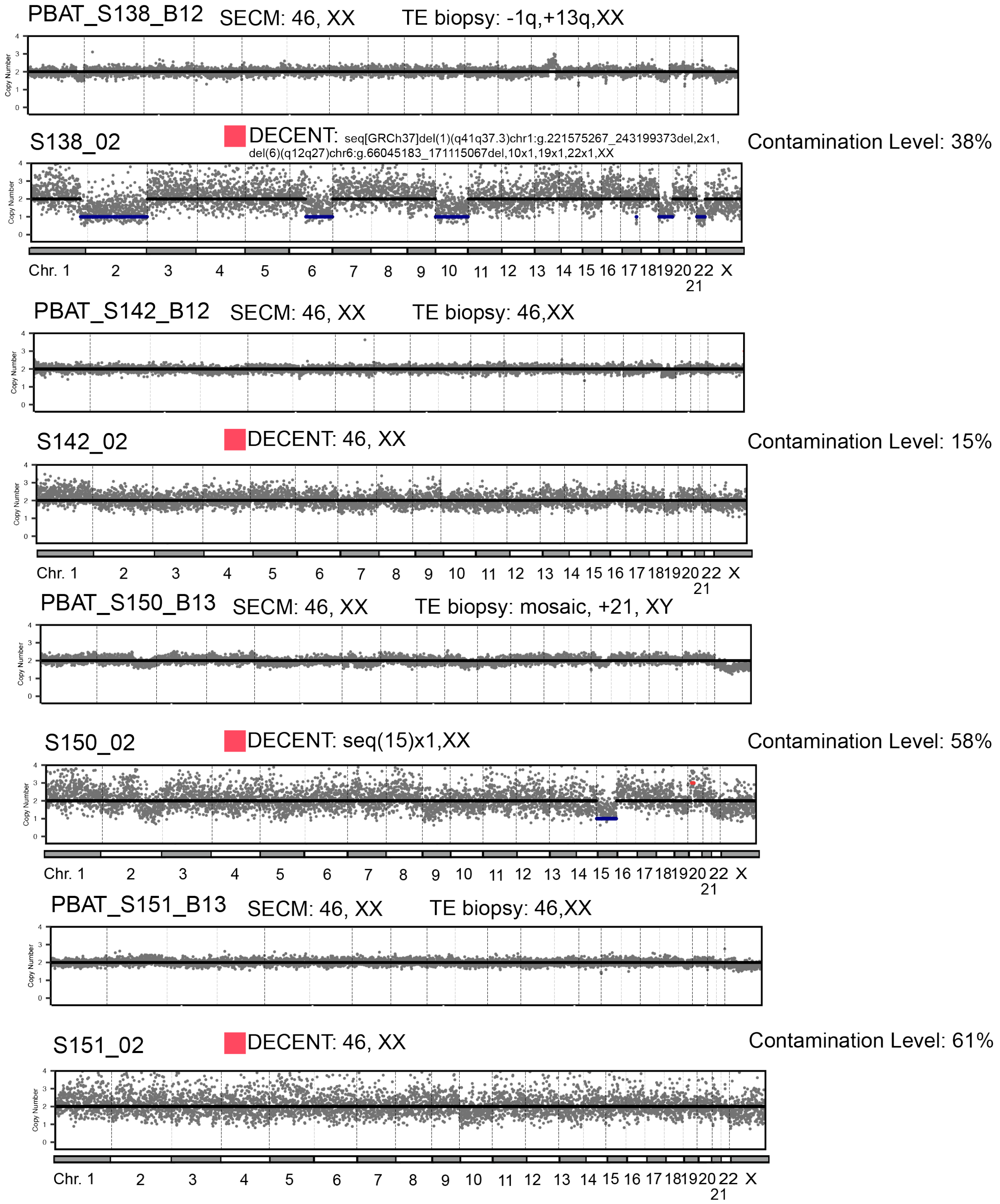

**Figure S18.** Analysis results of DECENT on moderate contaminated SECM samples (V). Each pair of plots represents a sample set, with the first plot displaying the chromosome copy number results of the original sample and the second plot showing the chromosome copy number results after applying DECENT filtering with a threshold of 0.2. We show the original SECM, TE and the processed DECENT results.

##### A.19 Supplementary Figure S19: Analysis Results of DECENT on Moderate Contaminated SECM samples (VI)

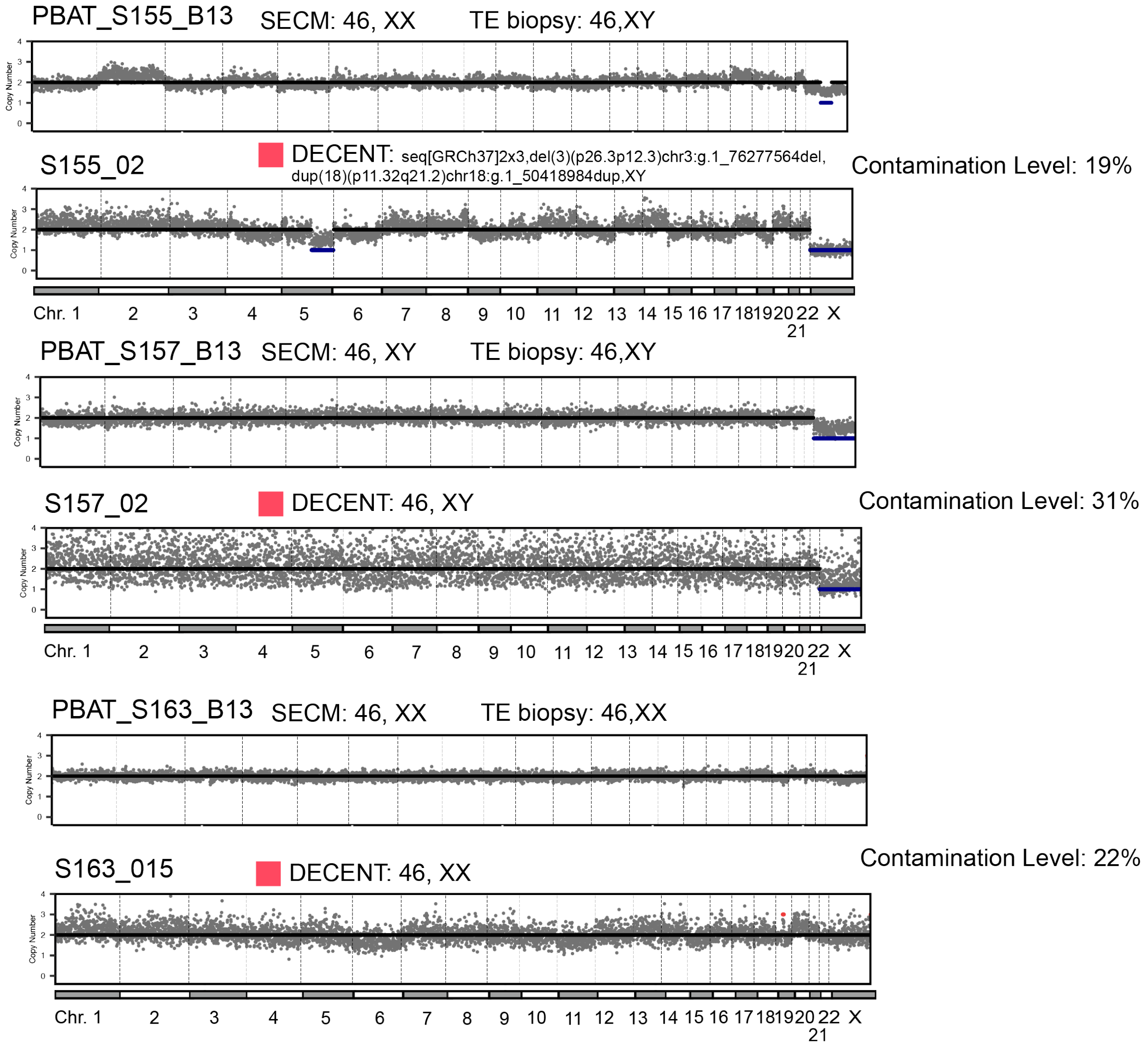

**Figure S19.** Analysis results of DECENT on moderate contaminated SECM samples (VI). Each pair of plots represents a sample set, with the first plot displaying the chromosome copy number results of the original sample and the second plot showing the chromosome copy number results after applying DECENT filtering with a threshold of 0.2. We show the original SECM, TE and the processed DECENT results.

#### A.20 Supplementary Figure S20: Analysis Results of DECENT on Moderate Contaminated SECM samples (VII)

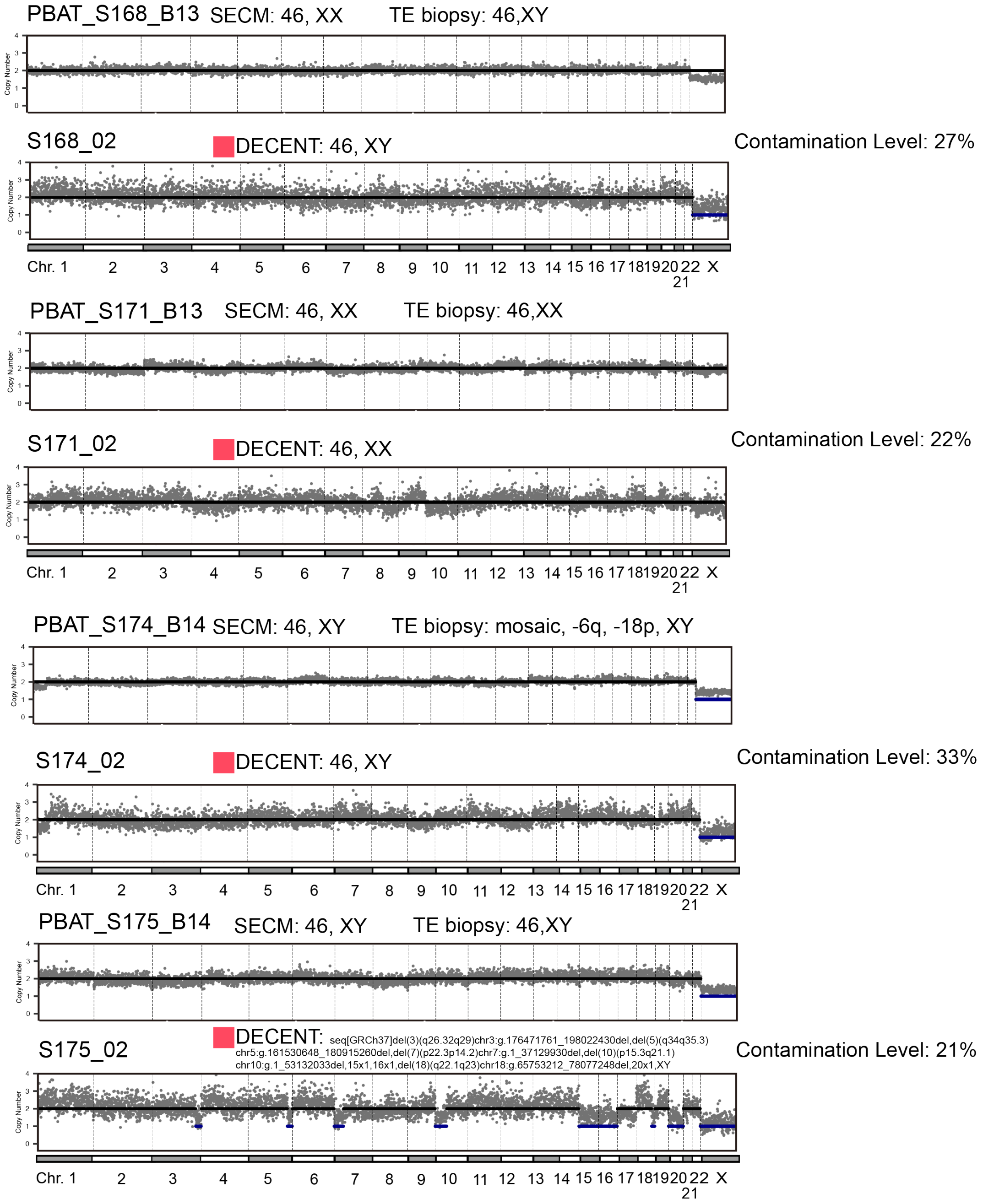

**Figure S20.** Analysis results of DECENT on moderate contaminated SECM samples (VII). Each pair of plots represents a sample set, with the first plot displaying the chromosome copy number results of the original sample and the second plot showing the chromosome copy number results after applying DECENT filtering with a threshold of 0.2. We show the original SECM, TE and the processed DECENT results.

##### A.21 Supplementary Figure S21: Analysis Results of DECENT on Moderate Contaminated SECM samples (VIII)

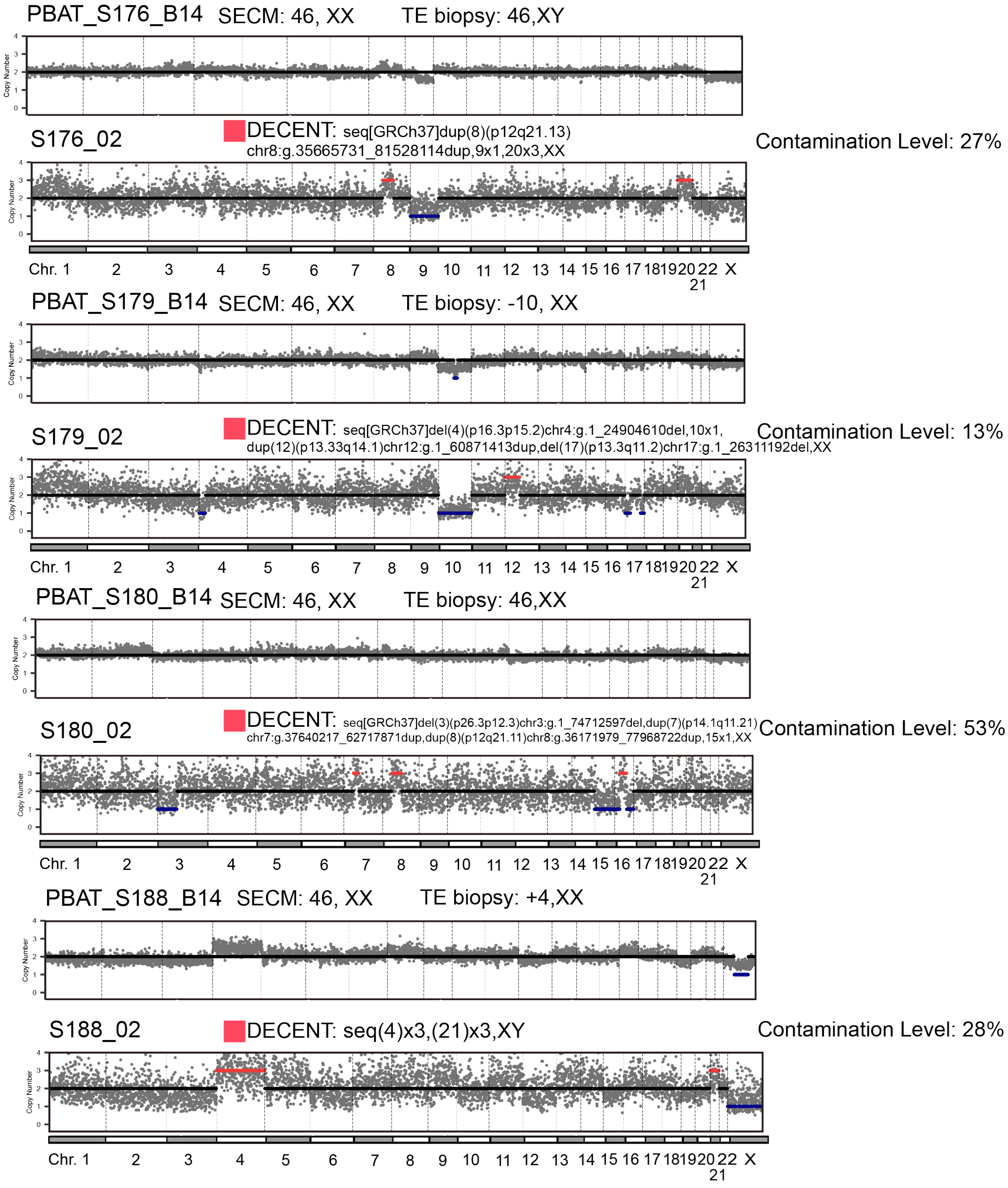

**Figure S21.** Analysis results of DECENT on moderate contaminated SECM samples (VIII). Each pair of plots represents a sample set, with the first plot displaying the chromosome copy number results of the original sample and the second plot showing the chromosome copy number results after applying DECENT filtering with a threshold of 0.2. We show the original SECM, TE and the processed DECENT results.

##### A.22 Supplementary Figure S22: Analysis Results of DECENT on Moderate Contaminated SECM samples (IX)

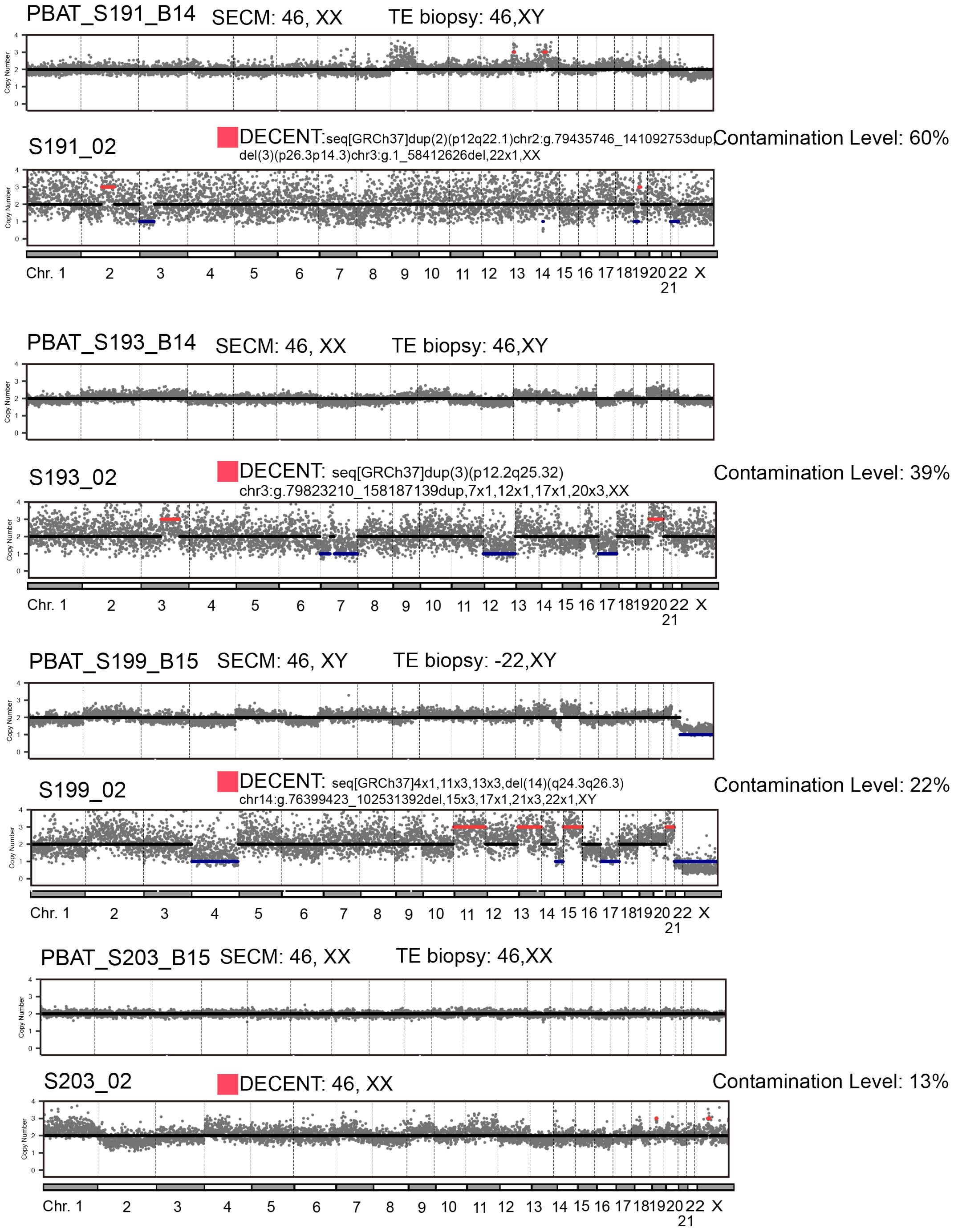

**Figure S22.** Analysis results of DECENT on moderate contaminated SECM samples (IX). Each pair of plots represents a sample set, with the first plot displaying the chromosome copy number results of the original sample and the second plot showing the chromosome copy number results after applying DECENT filtering with a threshold of 0.2. We show the original SECM, TE and the processed DECENT results.

##### A.23 Supplementary Figure S23: Analysis Results of DECENT on Moderate Contaminated SECM samples (X)

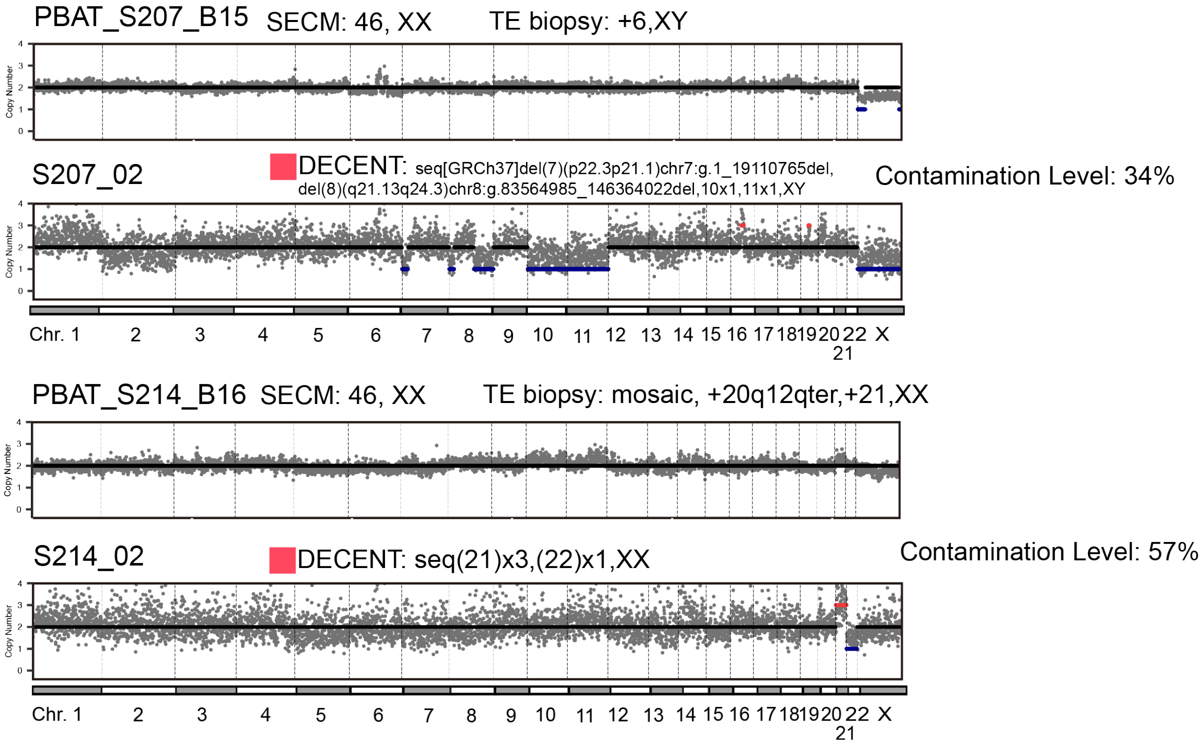

**Figure S23.** Analysis results of DECENT on moderate contaminated SECM samples (X). Each pair of plots represents a sample set, with the first plot displaying the chromosome copy number results of the original sample and the second plot showing the chromosome copy number results after applying DECENT filtering with a threshold of 0.2. We show the original SECM, TE and the processed DECENT results.

#### A.24 Supplementary Figure S24: Analysis Results of DECENT on Severe Contaminated SECM samples (I)

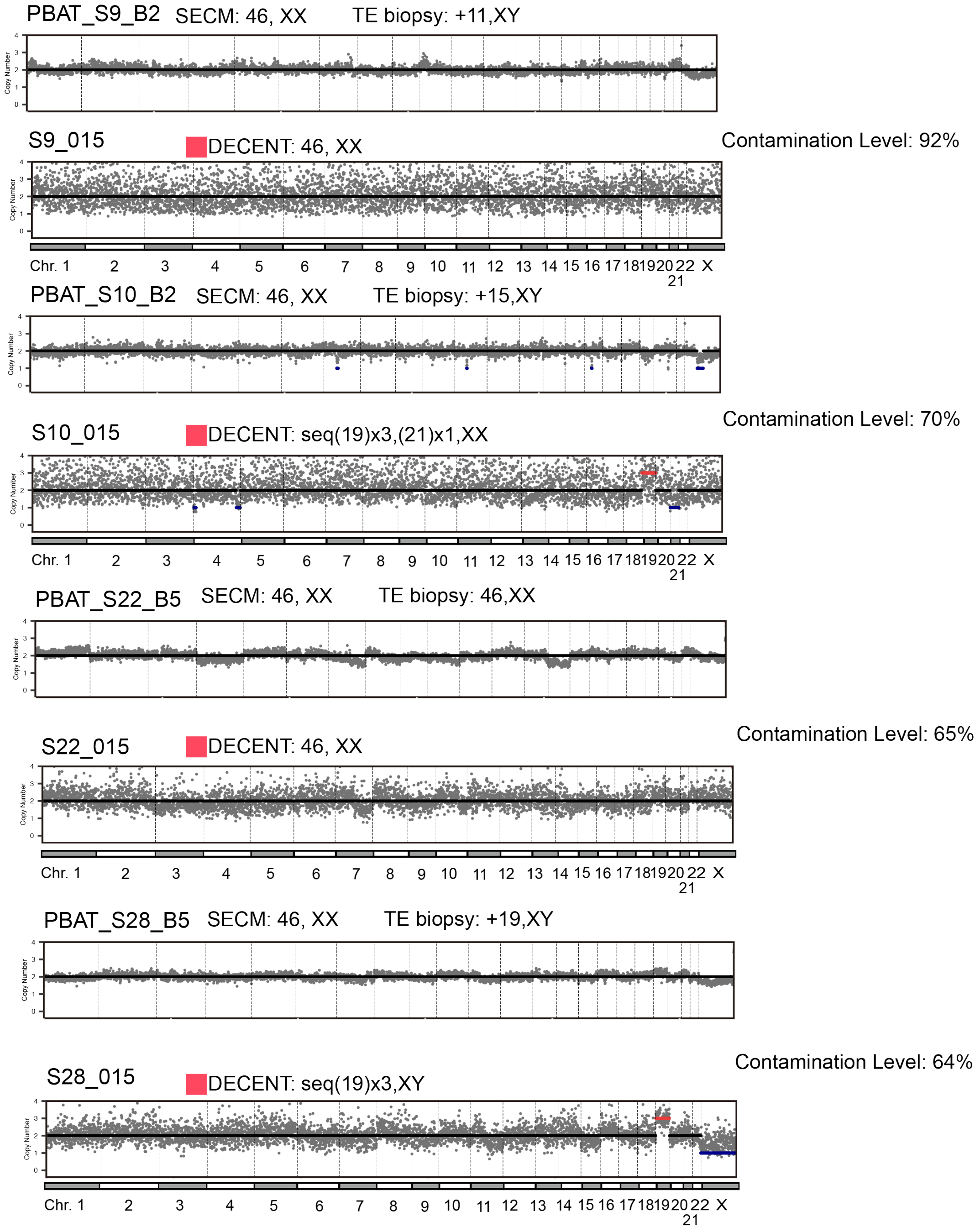

**Figure S24.** Analysis results of DECENT on severe contaminated SECM samples (I). Each pair of plots represents a sample set, with the first plot displaying the chromosome copy number results of the original sample and the second plot showing the chromosome copy number results after applying DECENT filtering with a threshold of 0.15. We show the original SECM, TE and the processed DECENT results.

#### A.25 Supplementary Figure S25: Analysis Results of DECENT on Severe Contaminated SECM samples (II)

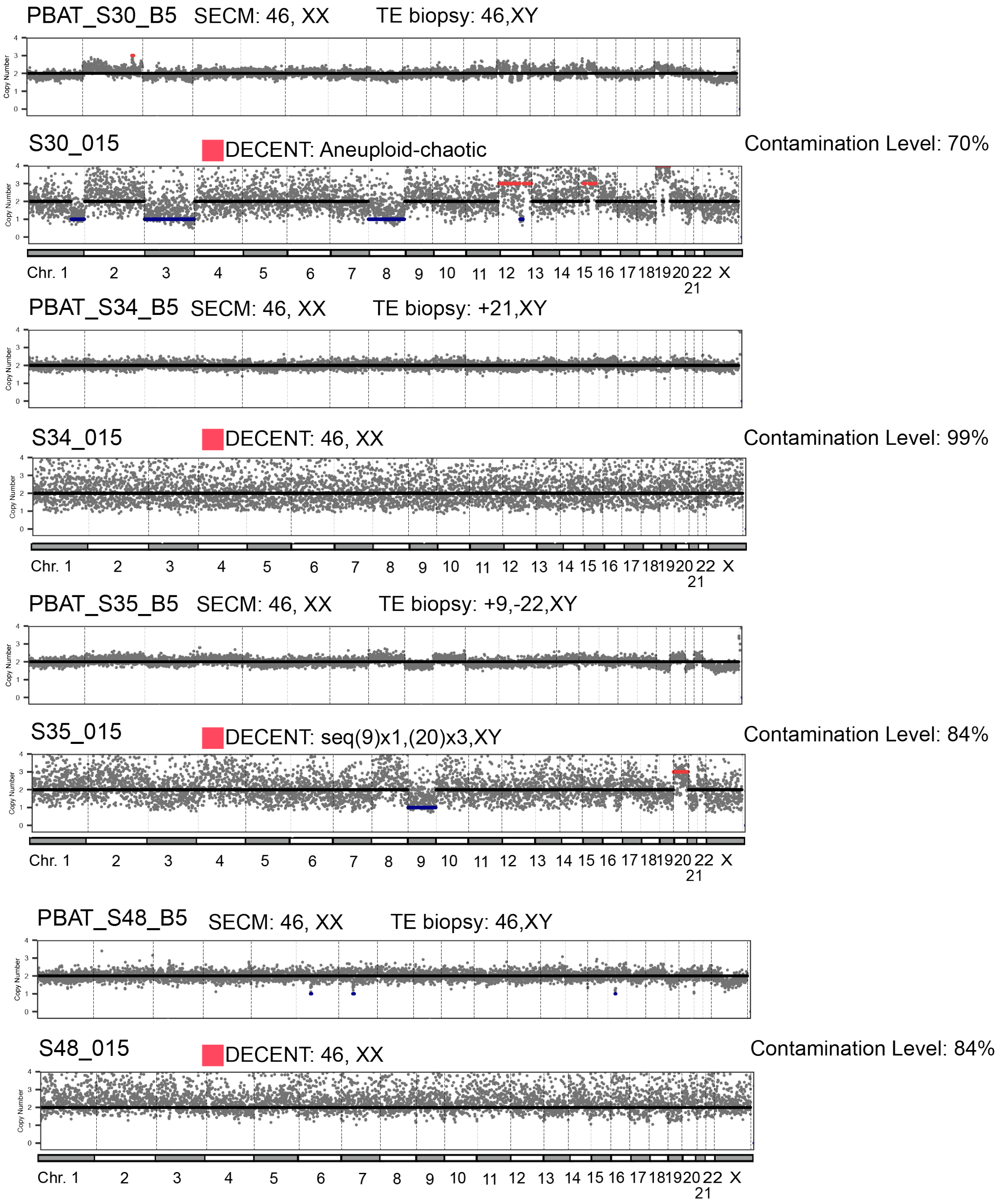

**Figure S25.** Analysis results of DECENT on severe contaminated SECM samples (II). Each pair of plots represents a sample set, with the first plot displaying the chromosome copy number results of the original sample and the second plot showing the chromosome copy number results after applying DECENT filtering with a threshold of 0.15. We show the original SECM, TE and the processed DECENT results.

#### A.26 Supplementary Figure S26: Analysis Results of DECENT on Severe Contaminated SECM samples (III)

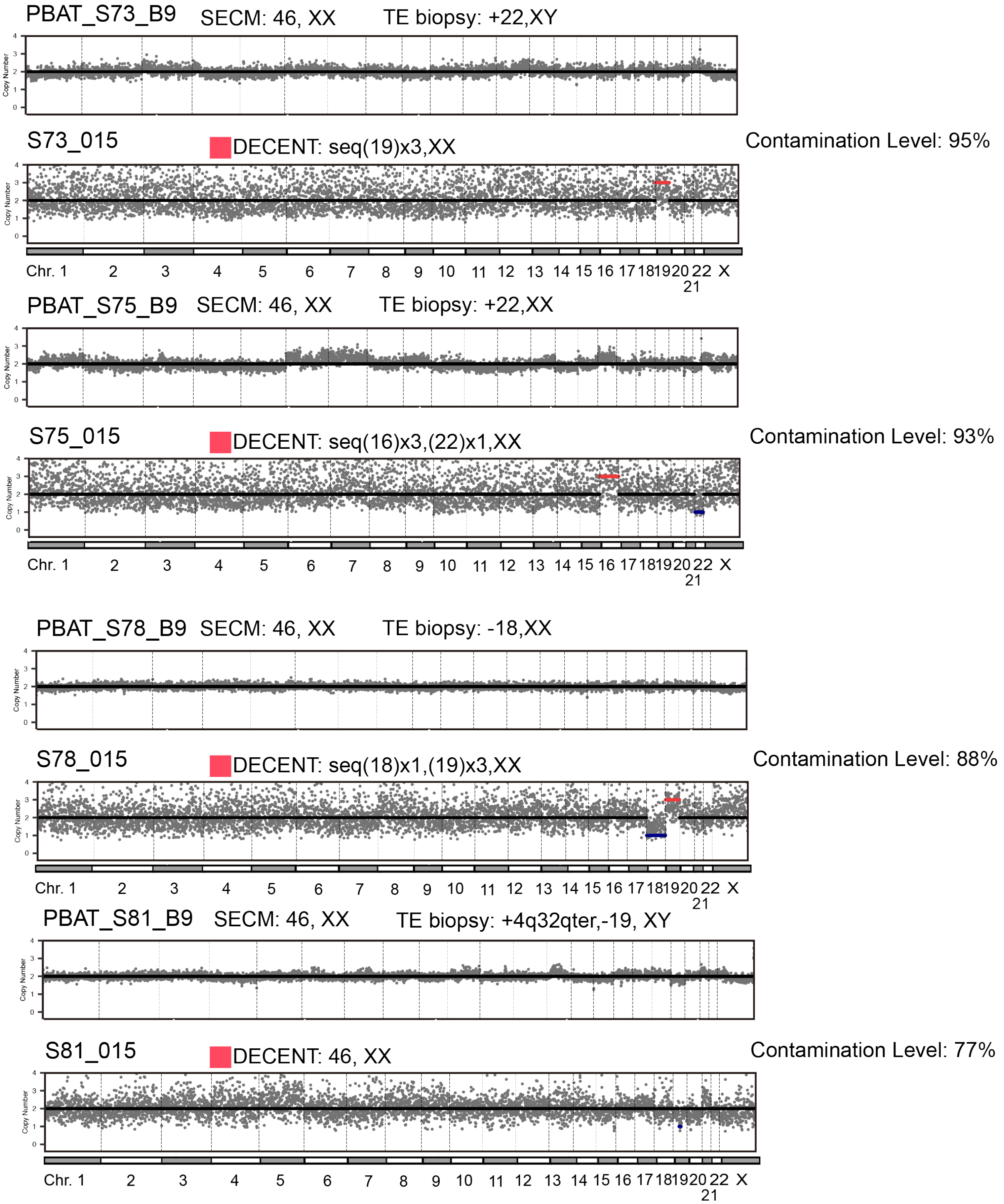

**Figure S26.** Analysis results of DECENT on severe contaminated SECM samples (III). Each pair of plots represents a sample set, with the first plot displaying the chromosome copy number results of the original sample and the second plot showing the chromosome copy number results after applying DECENT filtering with a threshold of 0.15. We show the original SECM, TE and the processed DECENT results.

#### A.27 Supplementary Figure S27: Analysis Results of DECENT on Severe Contaminated SECM samples (IV)

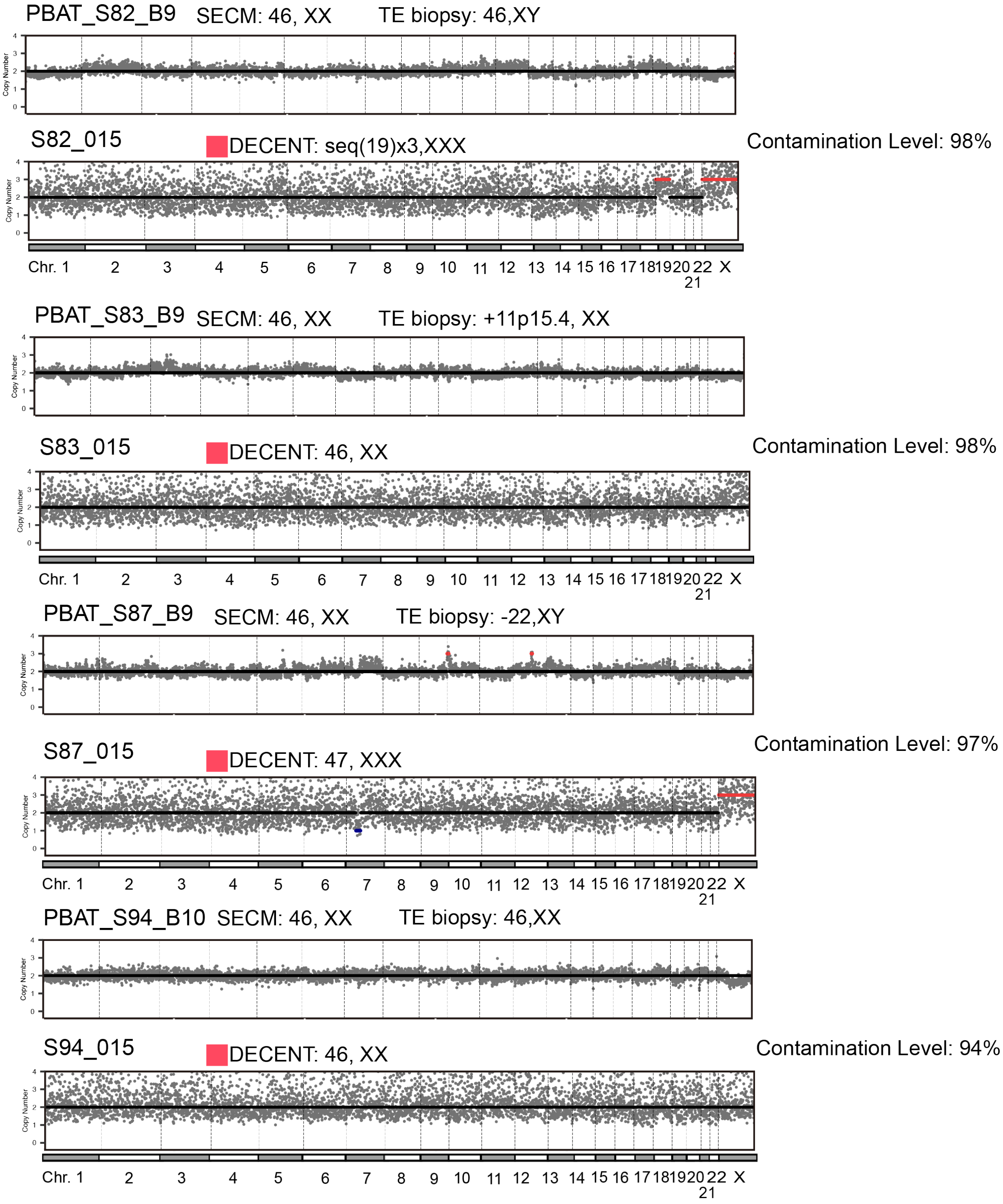

**Figure S27.** Analysis results of DECENT on severe contaminated SECM samples (IV). Each pair of plots represents a sample set, with the first plot displaying the chromosome copy number results of the original sample and the second plot showing the chromosome copy number results after applying DECENT filtering with a threshold of 0.15. We show the original SECM, TE and the processed DECENT results.

#### A.28 Supplementary Figure S28: Analysis Results of DECENT on Severe Contaminated SECM samples (V)

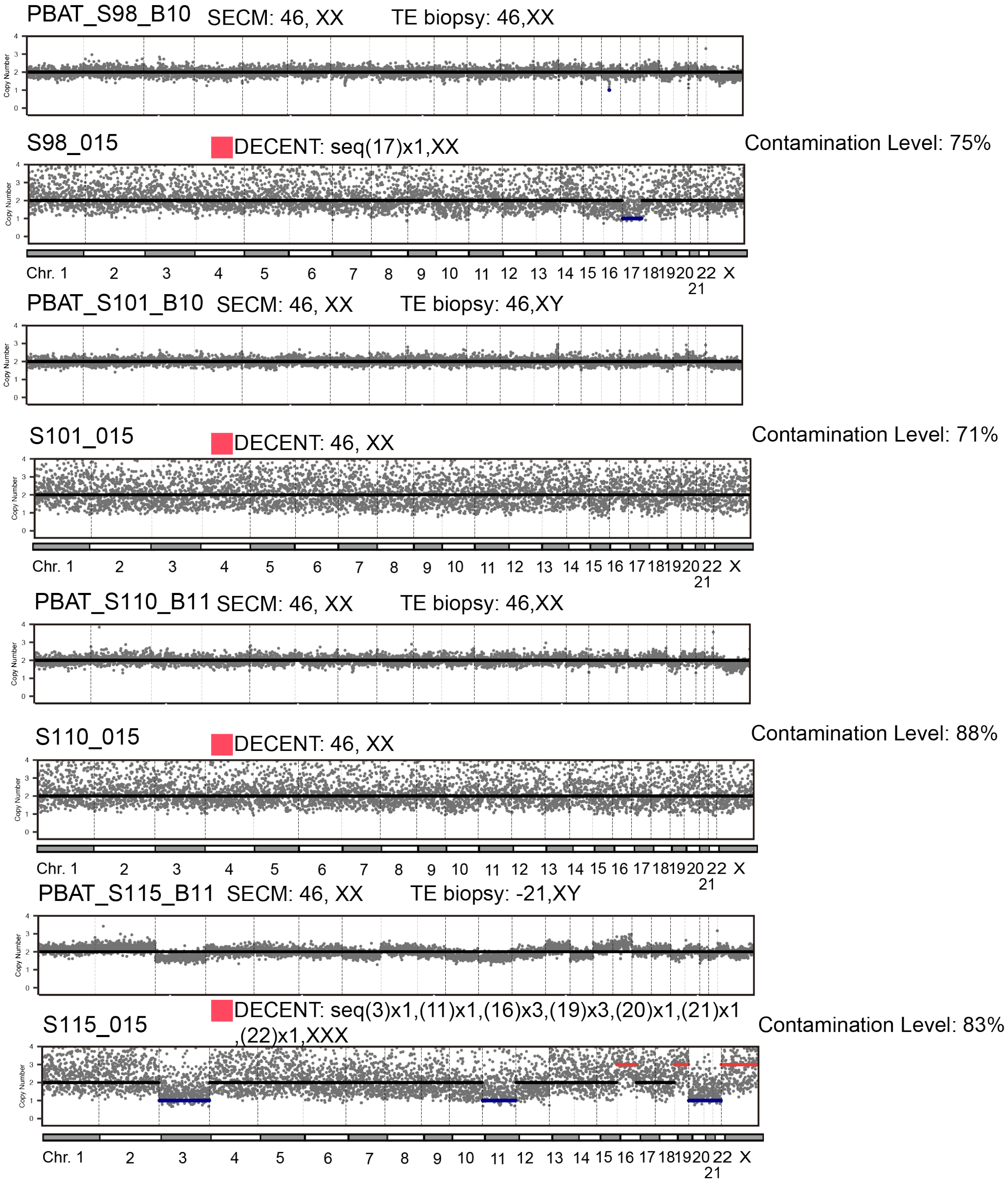

**Figure S28.** Analysis results of DECENT on severe contaminated SECM samples (V). Each pair of plots represents a sample set, with the first plot displaying the chromosome copy number results of the original sample and the second plot showing the chromosome copy number results after applying DECENT filtering with a threshold of 0.15. We show the original SECM, TE and the processed DECENT results.

#### A.29 Supplementary Figure S29: Analysis Results of DECENT on Severe Contaminated SECM samples (VI)

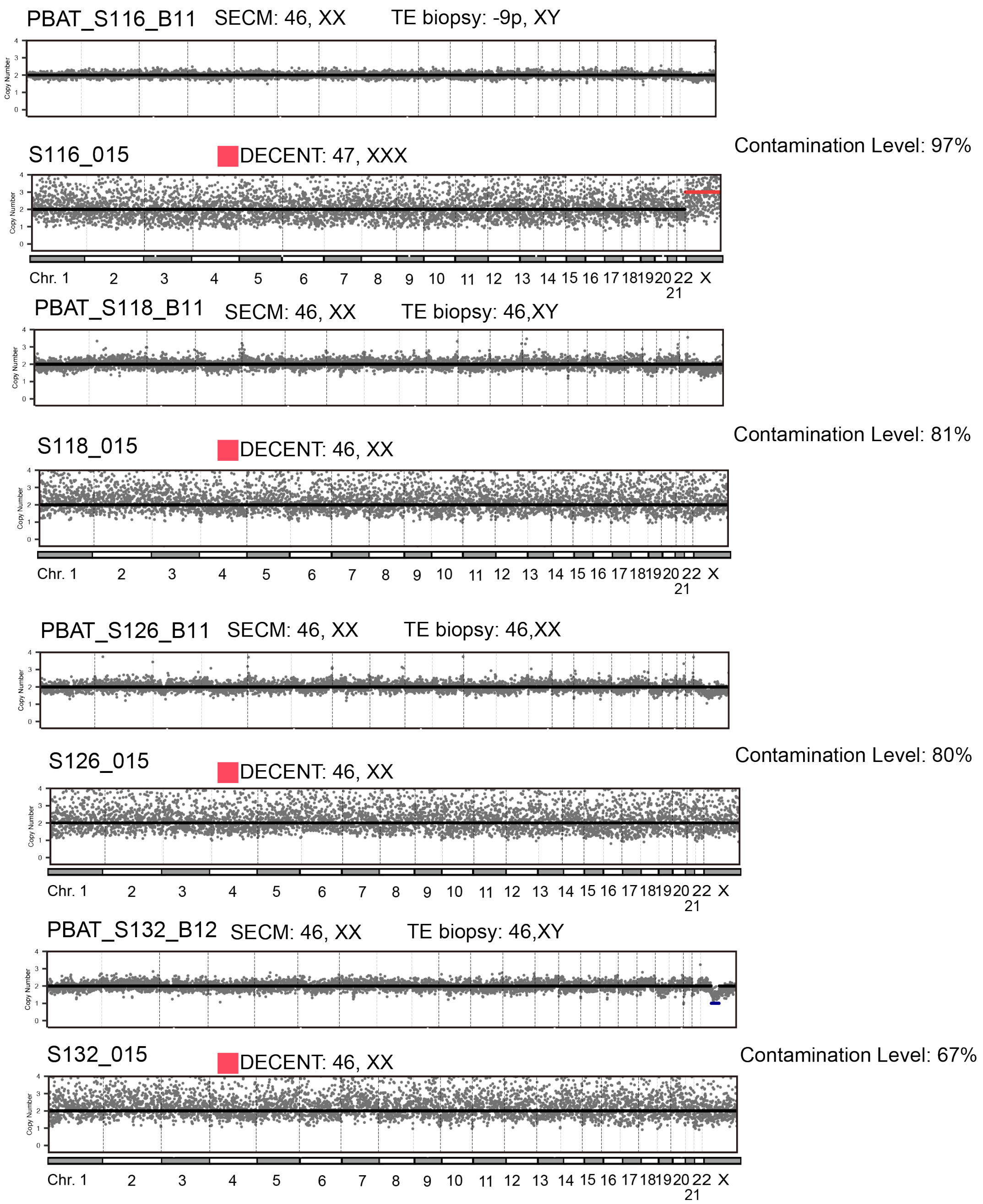

**Figure S29.** Analysis results of DECENT on severe contaminated SECM samples (VI). Each pair of plots represents a sample set, with the first plot displaying the chromosome copy number results of the original sample and the second plot showing the chromosome copy number results after applying DECENT filtering with a threshold of 0.15. We show the original SECM, TE and the processed DECENT results.

#### A.30 Supplementary Figure S30: Analysis Results of DECENT on Severe Contaminated SECM samples (VII)

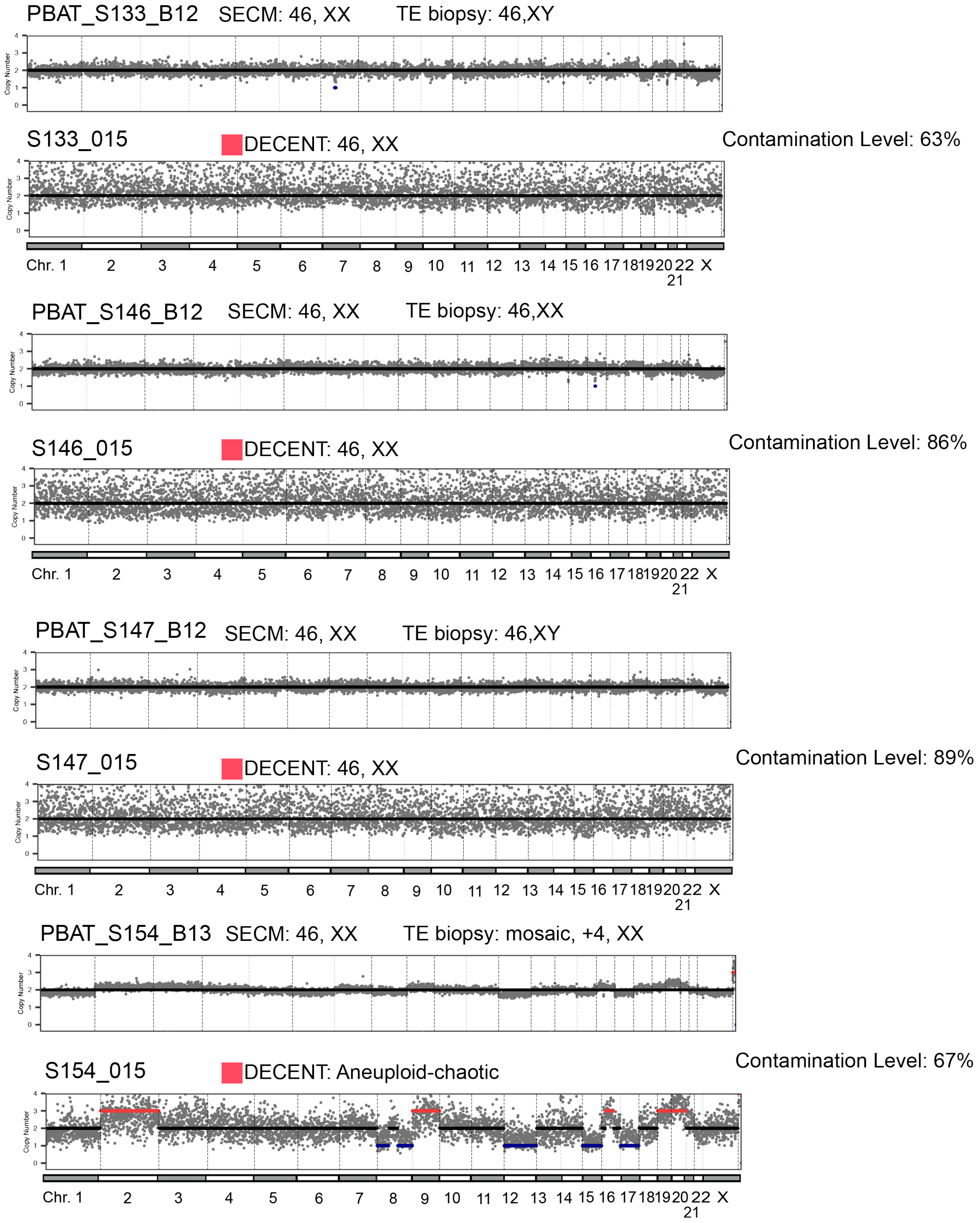

**Figure S30.** Analysis results of DECENT on severe contaminated SECM samples (VII). Each pair of plots represents a sample set, with the first plot displaying the chromosome copy number results of the original sample and the second plot showing the chromosome copy number results after applying DECENT filtering with a threshold of 0.15. We show the original SECM, TE and the processed DECENT results.

#### A.31 Supplementary Figure S31: Analysis Results of DECENT on Severe Contaminated SECM samples (VIII)

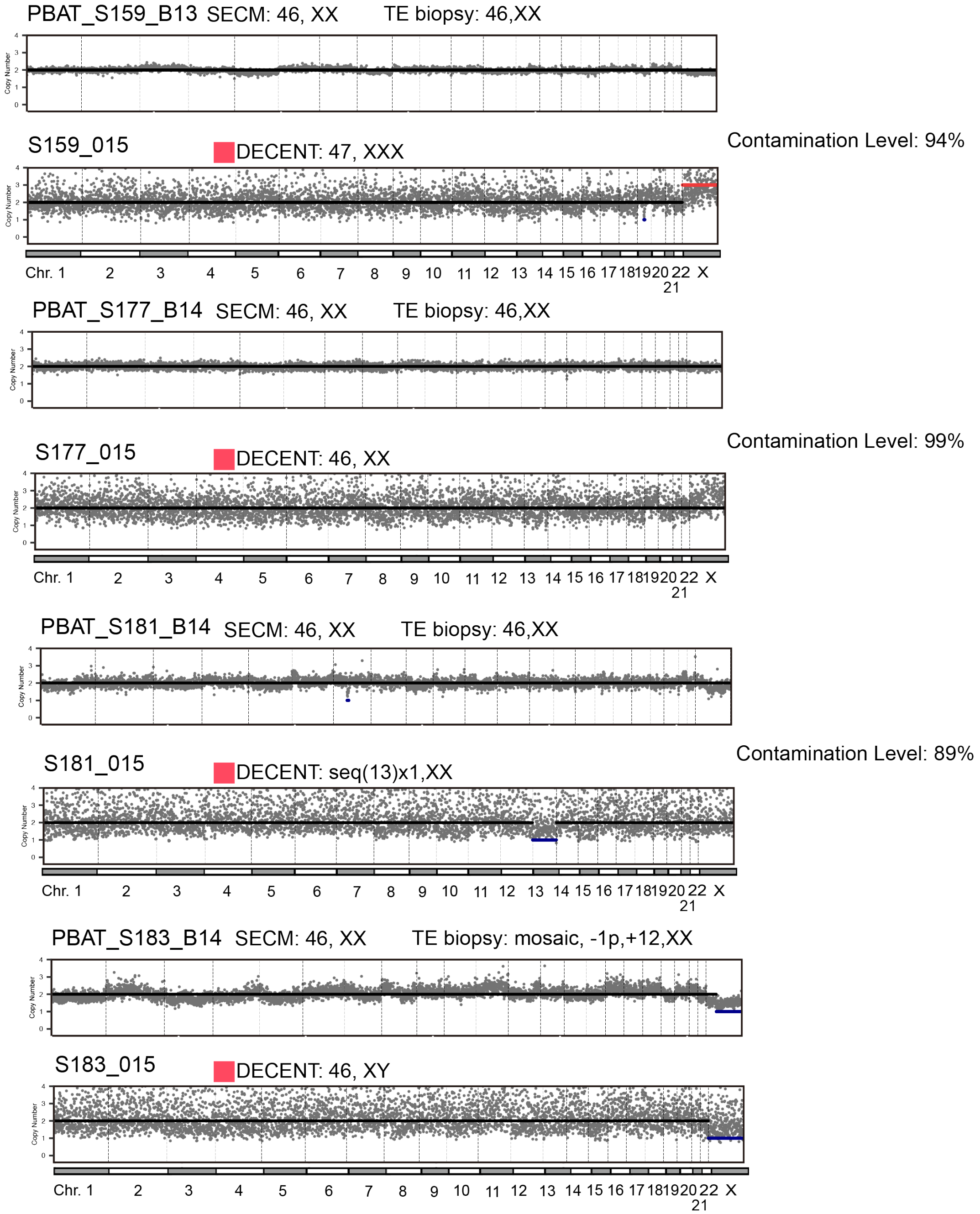

**Figure S31.** Analysis results of DECENT on severe contaminated SECM samples (VIII). Each pair of plots represents a sample set, with the first plot displaying the chromosome copy number results of the original sample and the second plot showing the chromosome copy number results after applying DECENT filtering with a threshold of 0.15. We show the original SECM, TE and the processed DECENT results.

#### A.32 Supplementary Figure S32: Analysis Results of DECENT on Severe Contaminated SECM samples (IX)

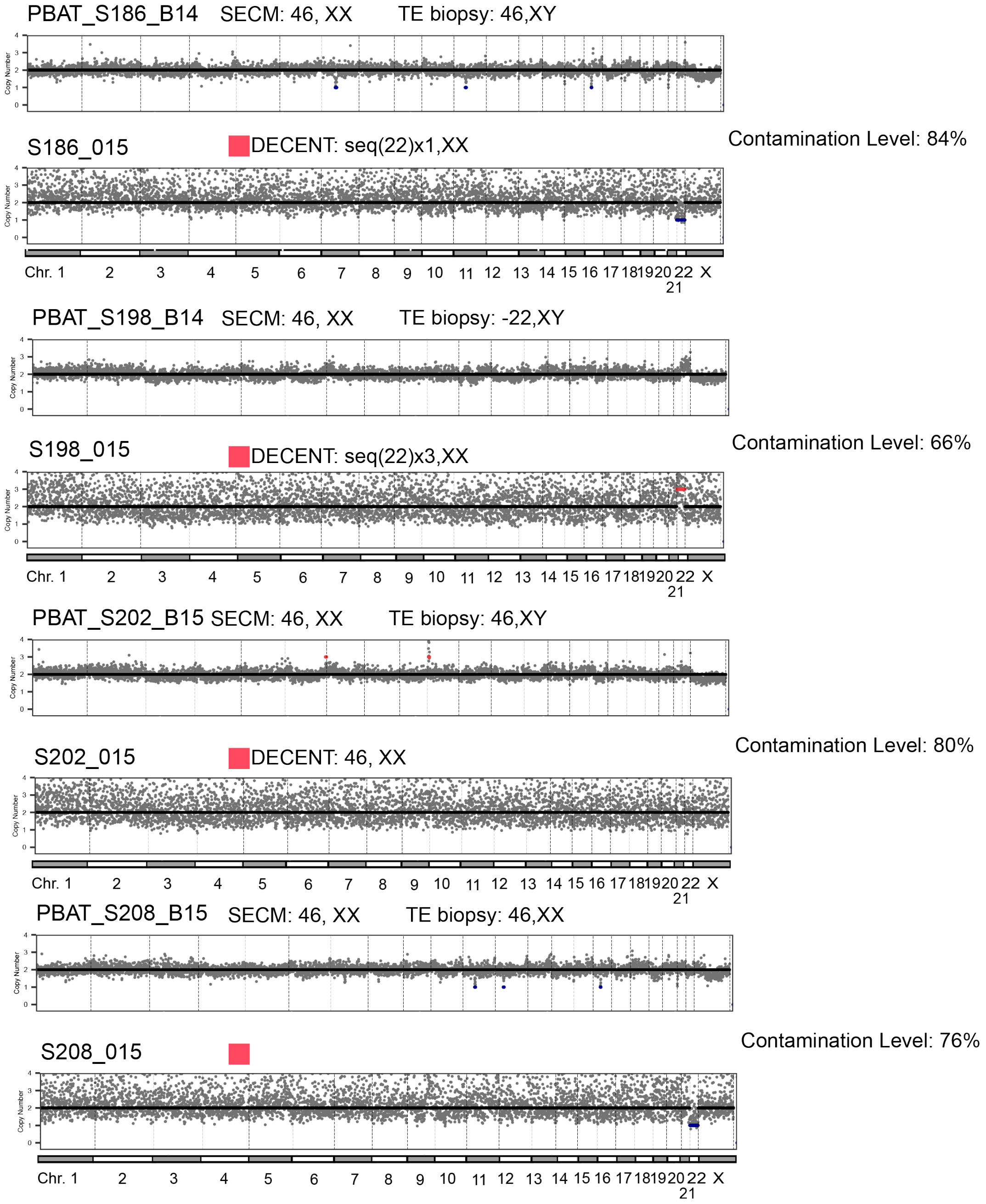

**Figure S32.** Analysis results of DECENT on severe contaminated SECM samples (IX). Each pair of plots represents a sample set, with the first plot displaying the chromosome copy number results of the original sample and the second plot showing the chromosome copy number results after applying DECENT filtering with a threshold of 0.15. We show the original SECM, TE and the processed DECENT results.

#### A.33 Supplementary Figure S33: Analysis Results of DECENT on Severe Contaminated SECM samples (X)

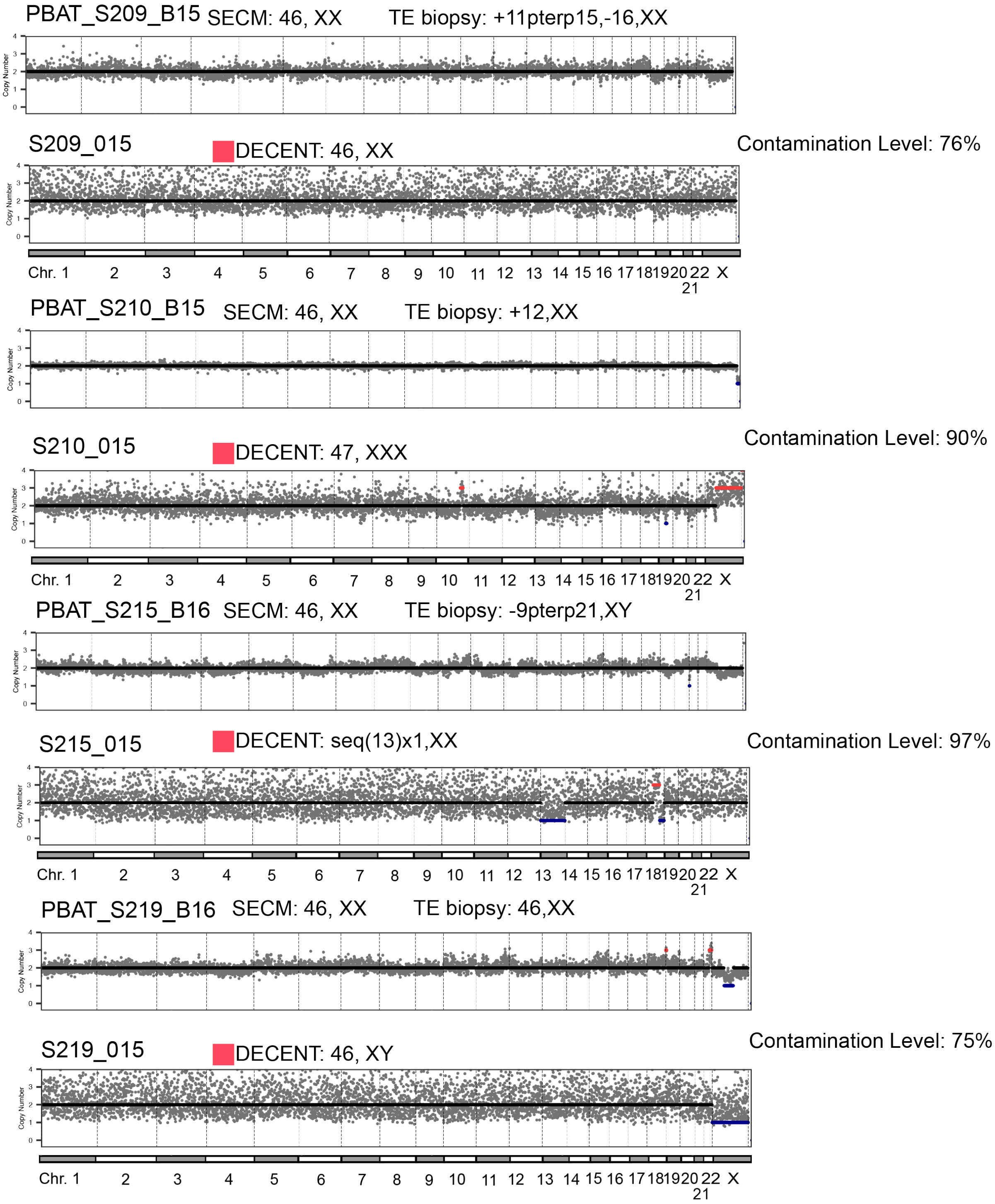

**Figure S33.** Analysis results of DECENT on severe contaminated SECM samples (X). Each pair of plots represents a sample set, with the first plot displaying the chromosome copy number results of the original sample and the second plot showing the chromosome copy number results after applying DECENT filtering with a threshold of 0.15. We show the original SECM, TE and the processed DECENT results.
